# Closed loop enhancement and neural decoding of human cognitive control

**DOI:** 10.1101/2020.04.24.059964

**Authors:** Ishita Basu, Ali Yousefi, Britni Crocker, Rina Zelmann, Angelique C Paulk, Noam Peled, Kristen K Ellard, Daniel S Weisholtz, G. Rees Cosgrove, Thilo Deckersbach, Uri T Eden, Emad N Eskandar, Darin D Dougherty, Sydney S Cash, Alik S Widge

## Abstract

Cognitive control is the ability to withhold a default, prepotent response in favor of a more adaptive choice. Control deficits are common across mental disorders, including depression, anxiety, and addiction. Thus, a method for improving cognitive control could be broadly useful in disorders with few effective treatments. Here, we demonstrate closed-loop enhancement of one aspect of cognitive control by direct brain stimulation in humans. We stimulated internal capsule/striatum in participants undergoing intracranial epilepsy monitoring as they performed a cognitive control/conflict task. Stimulation enhanced performance, with the strongest effects from dorsal capsule/striatum stimulation. We then developed a framework to detect control lapses and stimulate in response. This closed-loop approach produced larger behavioral changes than open-loop stimulation, with a slight improvement in performance change per unit of energy delivered. Finally, we decoded task performance directly from activity on a small number of electrodes, using features compatible with existing closed-loop brain implants. Our findings are proof of concept for a new approach to treating severe mental disorders, based on directly remediating underlying cognitive deficits.

## 1. INTRODUCTION

Mental disorders are a leading source of medical economic burden^1^. Current therapies do not target the cause of these disorders: dysfunctional communication in distributed brain circuits^2,3^. Electrical brain stimulation can effectively modulate such circuits^4^, but clinical neurostimulation trials in mental disorders have had mixed results^5,6^, for two reasons. First, spatially targeting a circuit may not be enough. Symptom-related activity is only occasionally present, and effective therapies may require temporal specificity: a closed feedback loop that detects pathological activity and intervenes to restore more efficient function^6–8^. Second, mental illness is heterogeneous. A label such as “depression” likely describes 10 or more different brain disorders^6,9^. Rather than trying to detect/treat ill-specified symptoms such as mood^10,11^, it may be more effective to target objective, rigorously measurable constructs, each representing a different component of cognitive/emotional processing^6,9,12^. In this dimensional approach, simultaneous impairments in multiple cognitive constructs act as “ingredients” to produce the clinical manifestation of disease^2,9^. An advantage of the dimensional approach is that individual constructs can be objectively measured, e.g. through performance on standardized tasks. That performance may be more easily reducible to specific brain circuits and stimulation targets.

One particularly promising construct is cognitive control, i.e. the ability to flexibly alter strategies/response styles as goals change^13–17^. First, cognitive control deficits are common across mental disorders, including depression, addiction, schizophrenia, and obsessive-compulsive disorder^18^. They are particularly relevant for mood and anxiety disorders, where patients are often unable to disengage from habitual, distress-driven behavior. Second, key aspects of control are readily measurable, using cognitive conflict tasks. Slower response times in these tasks signify difficulty exerting control to overcome response conflict, to the point that performance on such tasks is the most commonly used clinical and laboratory metric of cognitive control^18,19^. Third, control and conflict have a likely circuit substrate. Action selection in the face of competing options is well linked to interactions between multiple sub-regions of prefrontal cortex (PFC) and their counterparts in the striatum^20–22^. They similarly have a physiologic marker: conflict tasks evoke robust electrophysiologic signatures, namely theta (4-8 Hz) oscillations in PFC^23–25^ and low-frequency synchrony among frontal and subcortical structures^18,26–28^. Fourth, and most critically, that circuit substrate is well suited for manipulation through brain stimulation. PFC, striatum, and basal ganglia are densely interconnected by white matter tracts that run largely in the internal capsule^29,30^. Those tracts follow relatively stereotyped trajectories^29–32^, meaning that stimulation at a given capsular site will likely engage similar upstream/downstream circuitry across individuals. To that point, deep brain stimulation (DBS) of the internal capsule, DBS of subthalamic nucleus, and transcranial current stimulation of the lateral PFC all improve performance on cognitive control tasks^17,33,34^. In other words, the scientific basis exists for direct remediation of cognitive control deficits, if these disparate findings can be integrated.

Three large barriers have prevented that integration. First, there are no extant techniques for quantifying cognitive control performance in real time. Response time on conflict tasks, the standard metric, is also influenced by stochastic noise and often by changes in conflict level between subsequent task trials. Second, there are no proven ways to rapidly alter that performance, i.e. to respond to and cancel out momentary lapses. Third, although there are known physiologic signatures of cognitive control when averaged across time, there are no such signatures for moment-to-moment fluctuation in control. Here, we demonstrate methods to overcome those barriers, through a proof of concept in participants undergoing stereotaxic electrode monitoring for epilepsy (Figure 1A). We developed a method for quantifying conflict task performance at the single-trial level, while regressing out sources of undesired variability. This method is grounded in a state-space/latent-variable formalism, which generalizes the Kalman filtering approach that has been successful in other neural interface applications^35,36^. We showed that brief internal capsule stimulation rapidly enhances cognitive control. That enhancement was visible in raw task performance, in our latent variables, and in PFC theta oscillations. We then integrated those findings into a closed-loop controller that stimulated in response to lapses in cognitive control. Closed-loop stimulation produced larger performance changes than a corresponding open-loop paradigm. Finally, we showed that task performance, at the scale of single trials, can be decoded entirely from brain activity. Together, these findings provide proof of concept for a closed-loop system for treating cognitive control deficits.

**Figure 1:**
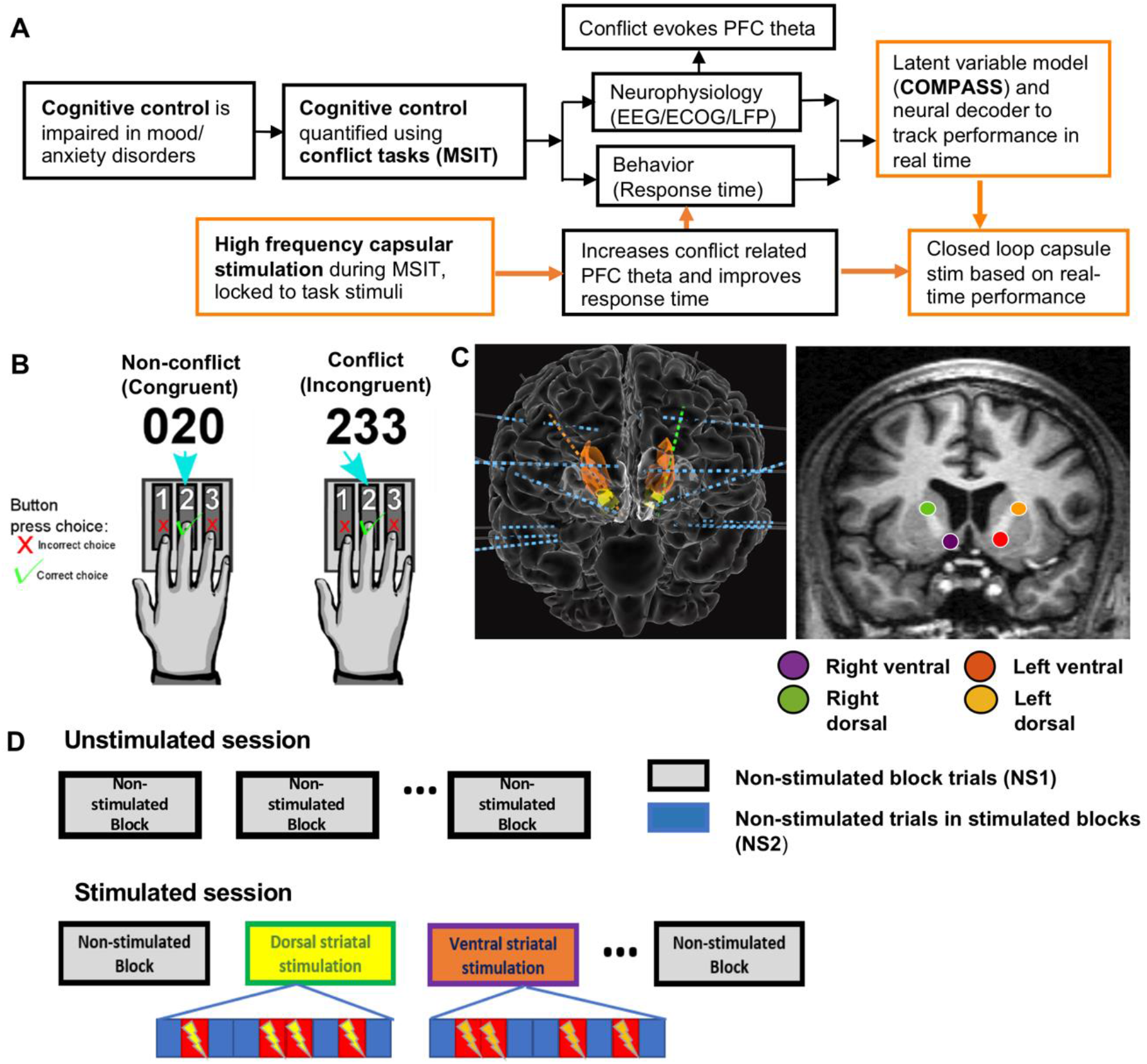
Experimental paradigms. A) Overview of cognitive conflict/control processing and the current study. Conflict, which requires high control, evokes PFC theta oscillations and slows decision (response) times. Orange boxes highlight this paper’s work. We developed intermittent stimulation approaches that improve these physiologic and behavioral correlates of control. We also developed a state-space model and neural decoder to track these quantities in real time. We then merged these for closed-loop enhancement of cognitive control, based on state-space behavior tracking. B) Schematic of the Multi Source Interference Task (MSIT), in which participants must inhibit pre-potent responses on 50% of trials. C) Schematic of a typical montage of depth electrodes (left) and stimulation targets (right). D) Trial structure during experimental sessions. In an unstimulated experimental session (top), blocks of trials were separated by brief rest periods. During an open-loop stimulation experiment (bottom), some blocks of MSIT trials had no stimulation at all (NS1), while others had stimulation only on a randomly-selected 50% of trials. Un-stimulated trials in these blocks are designated NS2.

## 2. RESULTS

### Cognitive control and theta power enhancement through internal capsule stimulation

Participants performed a cognitive control task (the Multi-Source Interference Task (MSIT), Figure 1B) while undergoing invasive multi-electrode monitoring. During task performance, we electrically stimulated in the internal capsule, at similar sites across participants (Figure 1C). We collected 8,790 trials across 176 blocks from 21 participants – 12 without brain stimulation, 5 with both unstimulated and open-loop stimulation sessions, 1 with only open-loop stimulation, and 3 with unstimulated and closed-loop stimulation sessions. Dropping incorrect responses and non-responses excluded 345 trials (3.92% of total; 8,445 trials retained in analysis). In open-loop experiments, a random 50% of trials received brief, task-linked stimulation (Figure 1D).

MSIT successfully engaged cognitive control: participants were 216 ms slower on high-conflict trials (N=21, p<0.001, t=33.62, Wald test on GLM coefficient, Figure 2A). Conflict increased task-related theta power in the posterior cingulate (p<0.02, t=3.24) and dorsolateral prefrontal cortex (p<0.001, t=4.31) (Figure 2B). Capsular stimulation enhanced both cognitive control and its electrophysiologic correlates, with dorsal sites showing stronger effects. Right dorsal (p<0.001, t=−4.28), left dorsal (p<0.01, t=−2.65) and right ventral (p<0.05, t=−2.64) capsular stimulation all significantly decreased reaction time (RT, Figure 2C, Table S3a) without impairing accuracy (Figure S2A). RTs under dorsal stimulation were faster than with ventral stimulation on both sides, with right dorsal being the overall most effective (Table S3a). These findings mirror the capsular topography, where more dorsal sites are enriched in fibers originating in dorsal PFC^1,2^, which in turn is associated with cognitive control during conflict tasks^1–4^. Consistent with that topography, stimulation in dorsal capsule sites outside of task performance propagated mainly to ACC and DLPFC (Figure S3). RT improvements were not explained by practice or regression to the mean; RT in the final non-stimulated (NS1) block was significantly higher than the initial block (Figure S4A).

**Figure 2:**
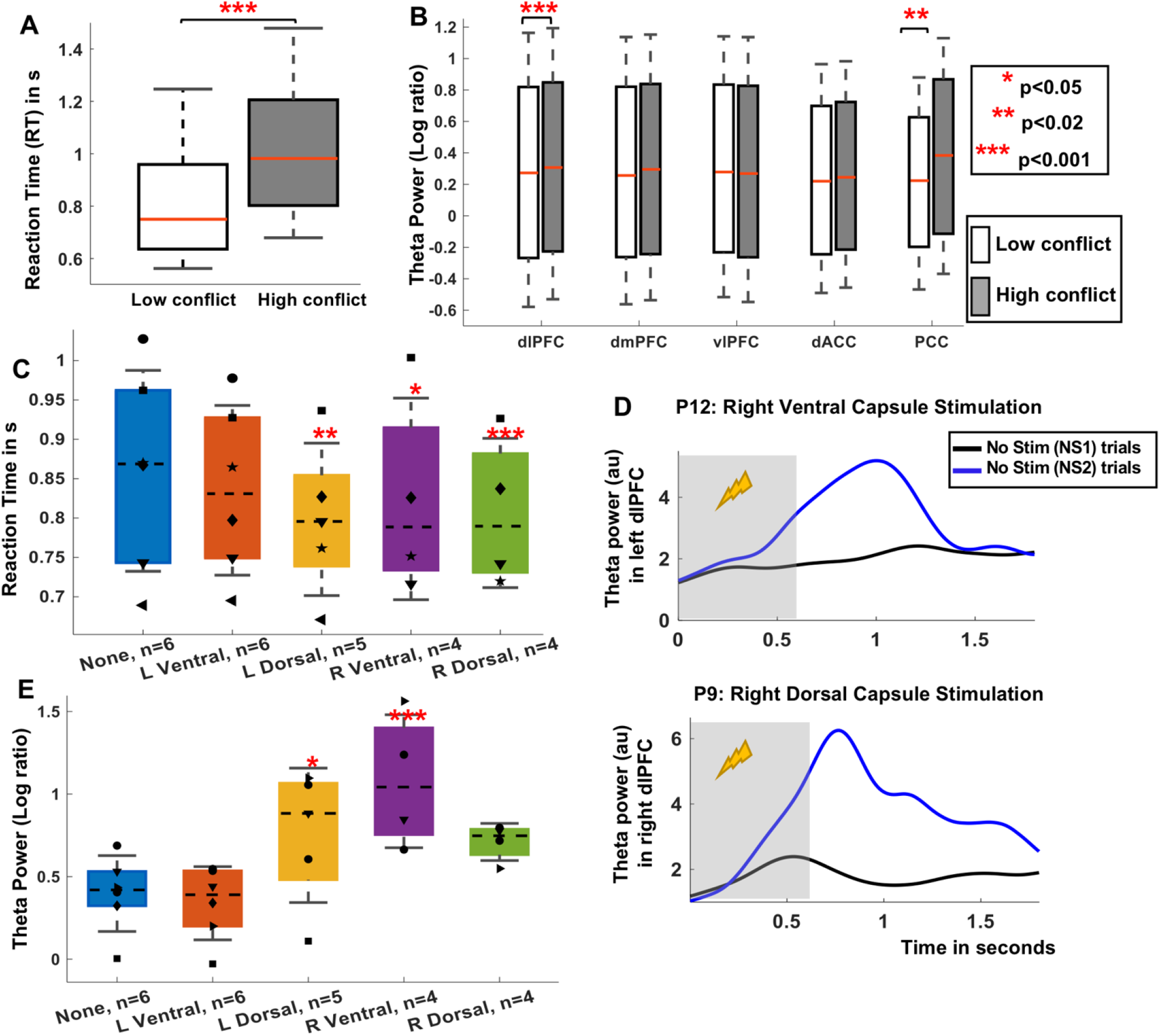
Effect of conflict and open-loop capsular stimulation on cognitive control: A) Reaction time (RT) during low- and high-conflict unstimulated trials (NS1) from 21 participants. B) Log theta power ratio during conflict and non-conflict NS1 trials in frontal regions. As in prior studies, conflict slowed responses and evoked significantly higher theta in dlPFC and PCC. C) RT during open loop stimulation during MSIT. Markers represent individual participants, bars show the mean, and error bars show standard error of the mean. Stimulation improved task performance, as reflected in lower RTs. Colors correspond to stimulation sites in Fig 1C, and inferential testing is performed against the unstimulated condition. D) Example theta power traces from PFC channels in 2 participants, with ventral (top) and dorsal (bottom) capsular stimulation. NS1 and NS2 trials are as in Fig 1D. The stimulation was from 0-0.6 seconds (grey window), although note that the curves show only data from non-stimulated trials. E) Task-evoked theta power (log ratio vs. pre-trial baseline), across PFC channels, in NS1 trials (None) compared to NS2 trials (all other conditions). Stimulation increased theta power, as in prior reports. In all panels, *, **, and *** denote p<0.05, 0.02, and 0.001 respectively, after appropriate correction. dlPFC, dorsolateral PFC; dmPFC, dorsomedial PFC; vlPFC, ventrolateral PFC; dACC, dorsal anterior cingulate cortex; PCC, posterior cingulate cortex.

There was no evidence for an interaction between stimulation and conflict level (AIC: −449.27 for a model without an interaction term vs. −445.72 with interaction). To assess stimulation’s effect on theta, we analyzed artifact-free trials interleaved within stimulated blocks (NS2, Figure 1D) and compared these to blocks without stimulation (NS1). Left dorsal and right ventral capsular stimulation significantly increased theta power in NS2 compared to NS1 trials (LD: p= 0.0428, RV: p=0.0006, FDR corrected, Figure 2D-E, Table S3b). Right dorsal capsular stimulation also increased theta but did not reach significance (p= 0.1733, FDR corrected). Theta increases were present in many PFC channels and were specific to behaviorally effective stimulation (Figure S5). Theta increases also could not be explained as regression to the mean or practice effects, as theta power attenuated over the experimental session in the absence of stimulation (Figure S4B).

### Closed-loop stimulation based on a state-space model efficiently enhances cognitive control

We next sought to quantify task performance and capsular stimulation effects at a trial-to-trial level. We achieved this with a state-space latent variable model (Figure 3A-B). This model assumes that each trial’s RT can be modeled as the combination of a baseline/expected RT for all trial types (*x*_*base*_), a specific slowing on high conflict trials (*x*_*conflict*_), and a Gaussian noise process. This formulation allows rapid tracking of stimulation-induced changes (see Methods). We verified that the two-process model captured the majority of variance in the RT data (Figure S6), that it converged in all participants (Figure S7), and that response accuracy did not carry additional information (Figure S8). Stimulation in the dorsal capsule improved overall performance (*x*_*base*_, Figure 3C) and reduced conflict effects (Figure 3D). Right dorsal capsular stimulation again had the largest effects. Ventral stimulation significantly reduced *x*_*conflict*_, but not *x*_*base*_. The observed differences could not be explained by block-to-block carry-over or other persistent effects, and *x*_*base*_ rapidly increased once stimulation ceased (Figure S9).

**Figure 3:**
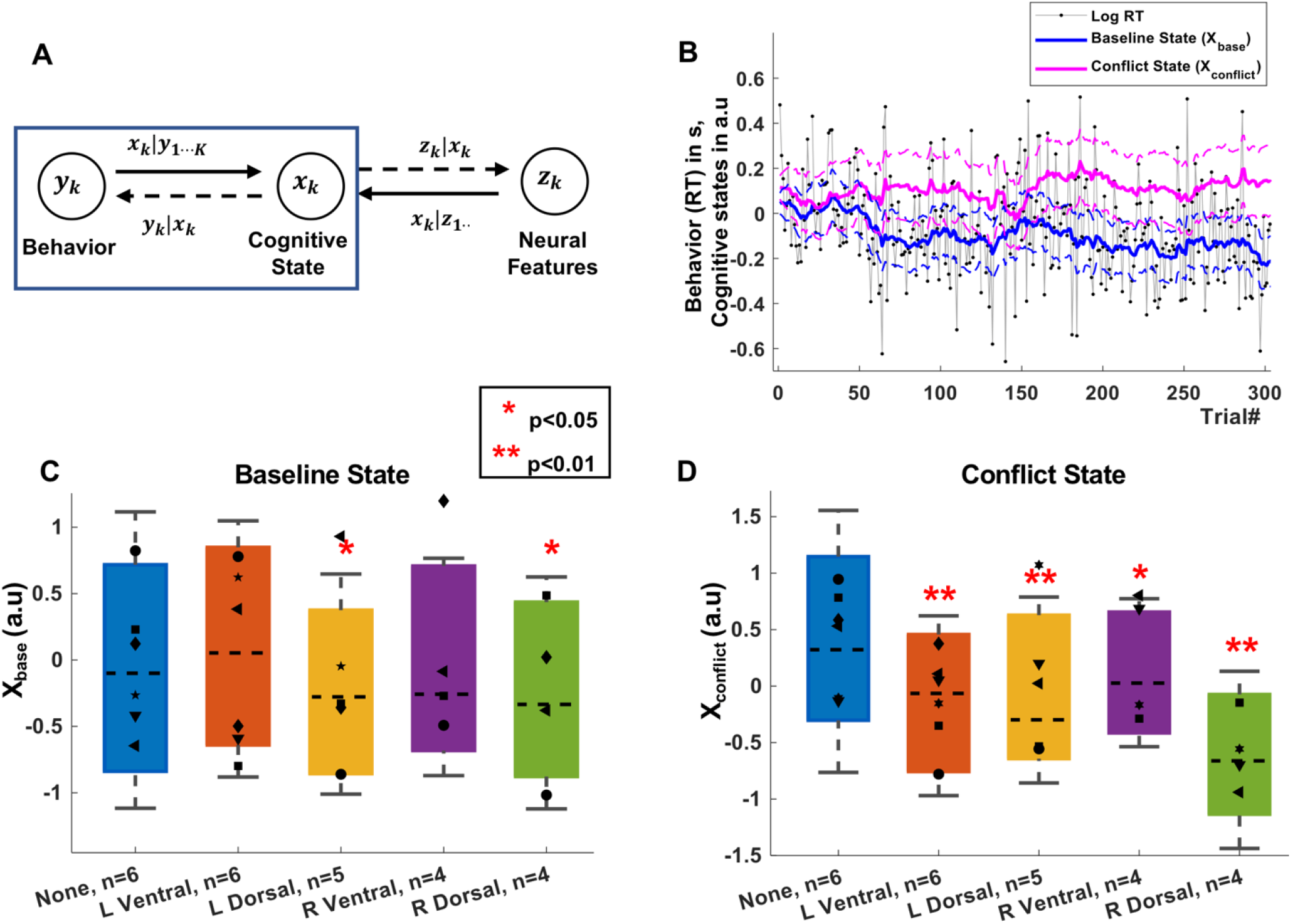
Effect of open-loop capsule stimulation on cognitive control: A) Schematic of the modeling framework, where behavior and neurophysiology are linked through a low-dimensional latent state space. Here, we focus on inferring latent states from behavior (blue box). B) Example of a participant’s raw behavior (RT) and its decomposition into *x*_*base*_ and *x*_*conflict*_. C) Effect of open-loop, randomly interleaved stimulation on *x*_*base*_ (expected RT). Only dorsal stimulation improved this task component. D) Effect of the same stimulation on *x*_*conflict*_ (expected RT to high conflict trials). Stimulation in all internal capsule sites altered this aspect of cognitive control, although right dorsal was still the most effective. Panels have the same formatting as Figure 2. * and ** represent p<0.05 and p<0.01 respectively, again corrected for multiple comparisons between stimulation and baseline. Statistical inference is through non-parametric permutations due to the highly autocorrelated nature of the state variables (see Methods).

We applied capsular stimulation under closed-loop control in 3 further participants. We estimated *x*_*base*_ in real time and triggered stimulation during control lapses, i.e. when *x*_*base*_ increased beyond an experimenter-determined threshold (Figure 4A). As predicted, conditioning stimulation on *x*_*base*_ specifically improved that variable (Figure 4B) without enhancing *x*_*conflict*_ (Figure 4C). Closed-loop stimulation was more effective than open-loop. Stimulation of the right ventral capsule, which did not have significant effects in open-loop tests (Figure 3C), now significantly reduced *x*_*base*_ (p<0.01, permutation test, Figure 4B). At both dorsal stimulation sites, closed-loop stimulation reduced *x*_*base*_ significantly more than open-loop stimulation (p<0.001, permutation test, Figure 4B). There were again no accuracy effects (Figure S2B). Closed-loop stimulation’s effect was manifest in raw RT data for right dorsal capsule stimulation (p<0.001, permutation test, Figure S10). The effects cannot be explained by regression to the mean (Figure S11). The closed-loop algorithm did stimulate more often that its open-loop counterpart (22-29 stimulated trials per block), but this alone did not explain the increased behavior effect. Closed-loop stimulation also appeared more efficient than open-loop in terms of performance gain for the applied energy. It produced a greater change in *x*_*base*_ per stimulation in the right ventral and dorsal capsule (Figure 4D), although this did not reach the pre-determined significance threshold (RV: p=0.207, RD: p=0.293).

**Figure 4:**
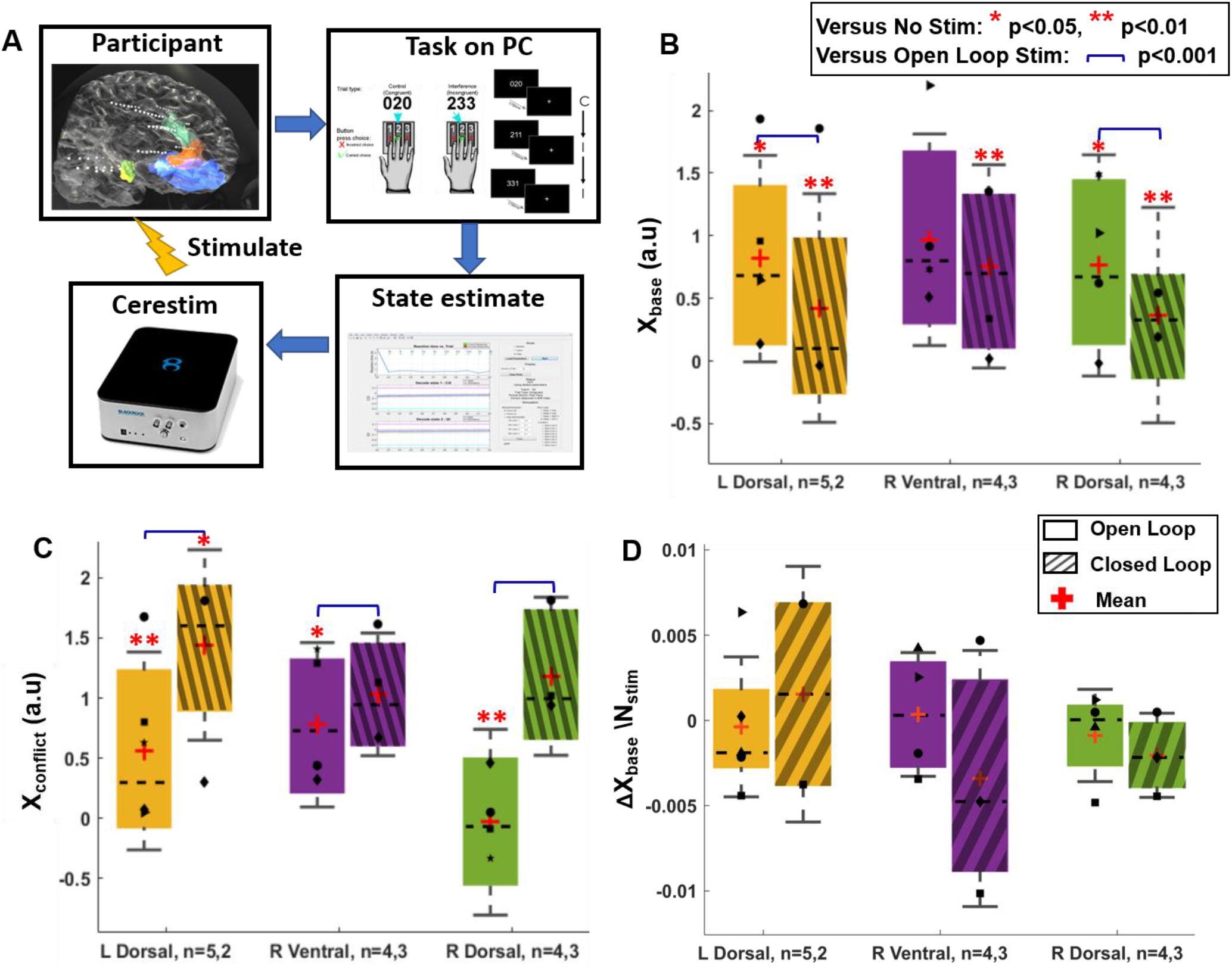
Closed-loop internal capsule stimulation efficiently enhances cognitive control. A) Schematic of closed loop paradigm. The controller estimated the Baseline state after each trial, commanding stimulation on the next trial if the state was above a pre-determined threshold. B) Effect of open- vs. closed-loop stimulation on *x*_*base*_. At multiple sites within the capsule, closed-loop stimulation was more effective at reducing *x*_*base*_ (improving task performance). C) Effect of open- vs. closed-loop stimulation on *x*_*conflict*_. Simulation conditioned on *x*_*base*_ did not reduce *x*_*conflict*_, and in fact significantly increased it at multiple sites. D) Comparison of open- vs. closed-loop effects on *x*_*base*_ (ΔXbase) (with 0 representing no change from NS1) divided by the number of stimulated trials (N_stim_). A negative value indicates a decrease (desired) in *x*_*base*_ caused by a specific stimulation on a block level. Panels B-D follow the same formatting as prior Figures. State values in B, C are normalized so that unstimulated blocks have a mean state value of 1 for each participant for both experiments, permitting comparison across participants. Significance in all panels is determined by a permutation test given the highly autocorrelated data. N for each experiment is given on the X-axis for open- and closed-loop participants.

A few participants reported that improving objective task performance also improved their subjective well-being. Although participants could not directly identify when stimulation was on/off, they perceived when they were performing more fluidly (Table S4). Two participants who had previously reported difficulty with effortful self-control noted relief of their usual anxiety (Table S5). No participant reported negative emotional effects during any stimulation experiment.

### Neural decoding of cognitive states for closed-loop control

To demonstrate that cognitive control lapses could be remediated outside of a controlled, structured task setting, we developed decoders to read out cognitive control from LFP. For each participant, we estimated an encoding model (Figure 5A) to map cognitive states to LFP power. State variables were linearly related to neural features (Figure S12), at a level exceeding chance performance (Figure S13). The confidence intervals of cognitive states decoded from LFP and estimated from behavior largely overlapped (Figure 5B, *x*_*base*_: 84.02 ± 15.8% overlap, *x*_*conflict*_: 83.17 ± 16.3% overlap). Decoding used relatively few power features in each participant (Figure 5C; 11.75 ± 6.63 features for *x*_*base*_ and 11.27 ± 6.74 for *x*_*conflict*_). Decoding weighted brain regions commonly implicated in cognitive control. *x*_*base*_ was encoded primarily in dlPFC (8-30, 130-200 Hz), vlPFC (15-30, 65-110 Hz), and temporal cortex (multiple bands). *x*_*conflict*_ was similarly encoded in dlPFC (most bands), vlPFC (8-55 Hz), temporal cortex (broadband), and amygdala (8-55 Hz).

**Figure 5:**
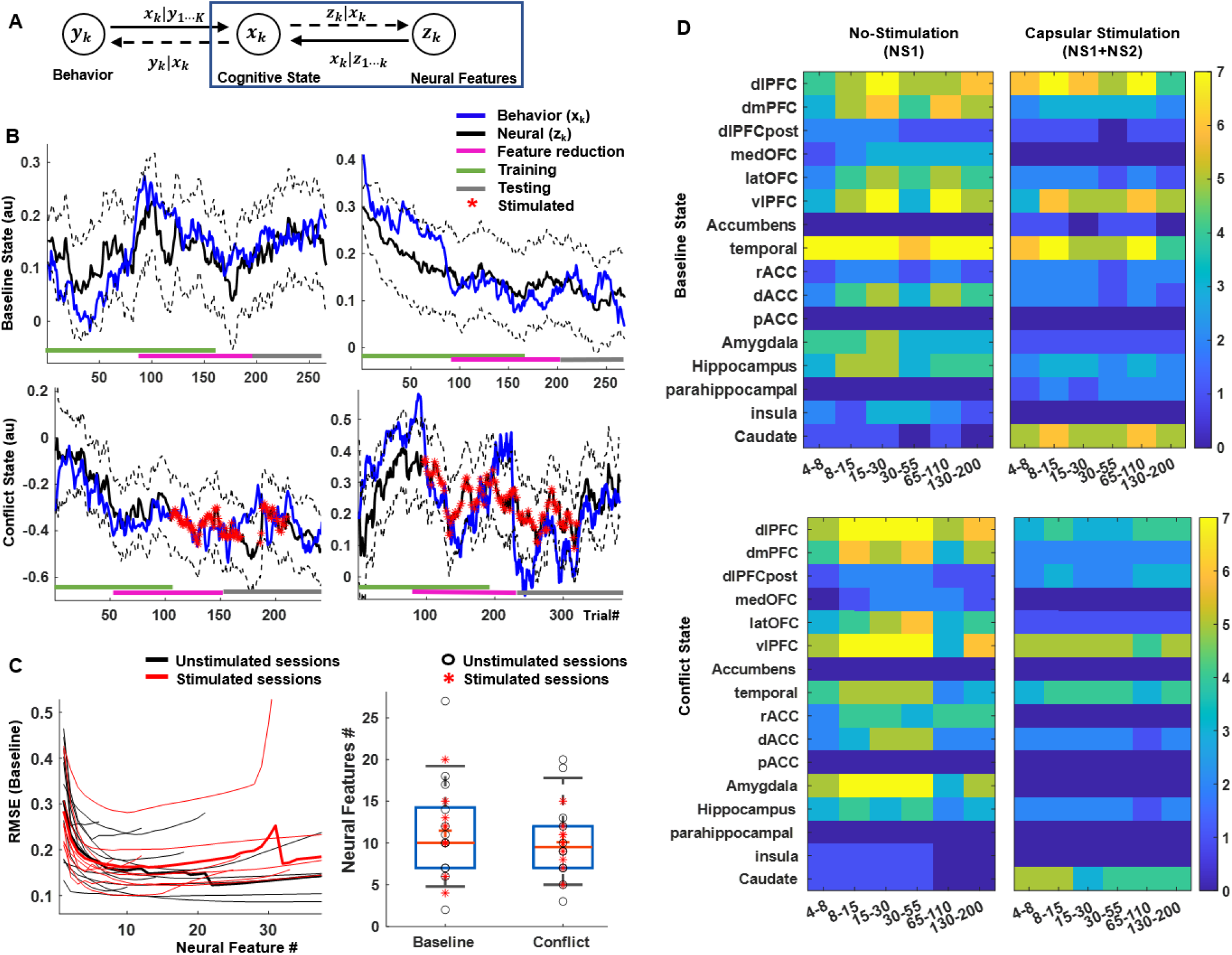
Neural decoding of cognitive states. A) Schematic of the encoding-decoding framework, which uses the same state variables as Figure 3. Here, we identify linear dependencies between neural features (LFP power) and the latent cognitive states (blue box). B) Examples, in two participants, of cognitive states as estimated from behavior and from neural decoding. Colored bars at the bottom of the plot indicate the data segments used for training, feature pruning (validation), and testing. There is strong agreement throughout the task run, including on data not used to train the decoding model. Decoding quality is similar during unstimulated (left) and stimulated (right) experiments. C) Optimum neural feature set determined using a feature dropping analysis (left). Thin lines represent individual participants, thick lines the mean. The solid circles indicate the number of features that minimizes root mean squared error (RMSE) for *x*_*base*_, for each participant, on a held-out validation set. This number of features did not differ between *x*_*base*_ and *x*_*conflict*_, or between stimulated and unstimulated blocks (right). D) Number of participants encoding *x*_*base*_ (top) and *x*_*conflict*_ (bottom) in different brain regions and frequency bands during non-stimulated (left, NS1) and intermittent capsular stimulation (right, NS1+NS2) blocks. Capsular stimulation shifts encoding from cortical regions to subcortical, particularly caudate.

Decoding was also possible during intermittent brain stimulation (*x*_*base*_: 83.15 ± 9.04%, *x*_*conflict*_: 84.43 ± 9.6% of trials overlapping the confidence interval of the behavioral estimate). Stimulation did not meaningfully alter the number of features needed for decoding (*x*_*base*_: 11.11 ± 4.73 vs. 11.75 ± 6.63 features; *x*_*conflict*_: 12.55 ± 4.24 vs. 11.27 ± 6.74 features; all p > 0.5, unpaired t-test). It did, however, decrease the number of cortical regions that encoded either *x*_*base*_ or *x*_*conflict*_ (Figure 5D). This encoding may have transferred to the dorsal striatum (caudate), which showed increased encoding across frequency bands, although this did not reach our pre-specified significance threshold for *x*_*base*_ (*x*_*base*_: t=−0.3980, p=0.6908, *x*_*conflict*_: t= 5.3947, p<0.001, paired t-test between encoder-model coefficients on NS1 and NS2 trials across participants).

## 3. DISCUSSION

Cognitive control is impaired in numerous mental disorders^6,12,18,37^. We augmented one aspect of human cognitive control, performance on a cognitive conflict task, by intermittent closed-loop stimulation of the internal capsule. The effects were detectable in both manifest data (raw reaction times) and derived variables. This brief intervention produced positive emotional effects in a small subset of patients who reported relevant pre-existing impairments. This may suggest a future utility of cognitive control as a psychiatric treatment target, independent of a patient’s diagnostic label. To that point, we previously showed that improvements in similar aspects of cognitive control may be a mechanism of action of clinical deep brain stimulation^17^. Further, we could separate and alter components of cognitive control. We enhanced baseline performance (expected RT on all trial types) without driving the conflict response (specific slowing and expected RT on high-conflict trials) in the same direction. This suggests that these two processes could be targeted separately, e.g. if only one was impaired in a hypothetical future patient. Both states could be decoded with a mean of 10 LFP spectral features per participant, from a mean of 6 brain regions. Existing human-grade implants may be able to implement these or similar decoders^1^.

Our result both converge with and diverge from past studies of cognitive control. Anatomically, we found the largest effects from stimulation in the dorsal capsule. This part of the capsule contains corticofugal axons originating mainly in lateral PFC and dorsomedial PFC/cingulate. We observed that lateral PFC in particular showed a theta power enhancement during behaviorally effective stimulation. Given the central role that those PFC regions likely play in cognitive control, the pattern we observed makes sense. We obtained our largest effects by stimulating in areas where we could modulate the relevant PFC sub-regions through retrograde activation. Our physiologic results support that retrograde model – effective stimulation increased PFC theta power, the most commonly reported correlate of control. Surprisingly, however, our decoders did not strongly select theta-band features when given the full range of LFP activity to consider. This may be because prior studies linking cognitive control to theta used EEG^23^ and were analyzed on a trial-averaged basis. The mean theta effect across trials, although robust, may not relate to the trial-to-trial performance. Further, LFP signals from a few millimeters of tissue may not correlate well with EEG, which records synchronized signals from broader swaths of cortex. Surprisingly, state variables were strongly encoded outside the canonical PFC-cingulate cognitive control network^18,37^, particularly in temporal lobe. Hippocampus and amygdala are important in human anxiety-like behavior^40^ and integrate signals from multiple prefrontal regions^41^. These structures may play a greater role in cognitive control than has been recognized.

Effective stimulation shifted encoding from fronto-temporal cortex to the dorsal striatum/caudate. This is consistent with models of cortico-striatal function that consider the dorso-medial striatum, including the caudate, to be part of a loop-structured circuit serving flexibility and exploration^18,20^. That encoding may also explain why dorsal capsular stimulation was more behaviorally effective, as the stimulation field likely overlapped both caudate and incoming fibers from dorsal PFC. It is less clear why stimulation slightly decreased the overall encoding of control-related variables, yet improved task performance. One possibility is the hypothesized link between cognitive control and anxiety^42^. Humans may normally exert more cognitive control than we need to, i.e. over-weighting risk. Our performance enhancement could be explained as a reduction of that “excess” control. This is concordant with the two participants who reported that stimulation allowed them to shift attention away from anxious, self-focused processing. We note that this is also a very different subjective effect than reports from clinical DBS of the capsule/striatum, which usually causes euphoria^43,44^. It seems unlikely that such positive mood changes drove faster RTs, because the majority of participants reported no subjective mood alteration.

Because participants had seizure vulnerability, we were limited to a single stimulation session between the completion of clinical monitoring and electrode removal. We could not retest the same stimulation multiple times in the same participant, nor could we attempt a variety of setpoints to determine the limits of this closed-loop approach. We were limited to intermittent stimulation to reduce seizure risk, and could not directly test this intermittent paradigm against the continuous DBS of prior studies. Electrodes were placed along individualized trajectories, and their location within the capsule varied substantially (Figure S14). In this context, we were likely able to find a consistent effect across participants because the topographic structure of the capsule is relatively consistent across individuals and species and is similarly consistent even when moving several millimeters rostro-caudally^29,30^. Electrodes used only for recording had a similar limitation. Demonstration of a consistent, repeatable within-subject effect would be critical for turning these results into a therapy. The nearly identical effects between this study and our prior work with psychiatric DBS patients^17^ do suggest that the effect is robust. Further, because there is known variability of fiber placement within the capsule despite the topographic map^31^, we might expect much larger effects in a clinical scenario where we could precisely optimize each electrode’s placement to engage a target of interest.

We enhanced *x*_*base*_, which reflects overall attentional focus. *x*_*conflict*_ corresponds to the more immediate effect of conflict and the difficulty of executing control. In a clinical setting, either might be disrupted, and we may need to apply closed-loop control to both simultaneously. Here, when we controlled *x*_*base*_, *x*_*conflict*_ significantly increased. These two states are not inherently anti-correlated, because both were reduced by open-loop stimulation (contrast Figure 3C-D against Figure 4B-C), but it remains to be shown that *x*_*base*_ can be altered without effects on other cognitive variables. Similarly, our two-state model follows one specific theory, where control is allocated reactively in response to decision conflict. There are other theoretical models of cognitive control measurement and allocation^37^, and it would be important to explore how our stimulation approaches do or do not alter variables in those frameworks. For instance, alternate frameworks place a heavy emphasis on a participant’s motivation to exert control, sometimes framed as expected value. Our results could be explained by an increase in that expected value, and dissecting such questions will require more sophisticated tasks. Finally, decoding might perform better on network-level measures rather than the LFP power features we used, based on recent studies linking connectivity metrics to cognitive control^18,26–28^. At present, connectivity is difficult to estimate in real time or on a therapeutic device, although approaches are emerging^45^.

Translating these findings to practical clinical use will require overcoming both technical and ethical hurdles. Technically, there is substantial interest in the idea of cognitive control as a cross-cutting symptom of numerous psychiatric disorders^18,38,46^, and thought leaders in psychiatry are urging the development of treatments that directly target decisional/cognitive dysfunctions^47,48^. Another group recently reported closed-loop control of impulsivity, a close cousin of cognitive control^49^. We and others have argued that cognitive remediation is a mechanism of multiple existing neurostimulation therapies, but is not recognized as such^6,34,50^. There are not, however, well-established diagnostic and monitoring scales for cognitive control or most other cognitive constructs. There are promising first steps^38,51^, but establishing clinically valid assessments would be important. It would similarly be both important and interesting to understand how these results might change with a different task that assesses different aspects of cognitive control, e.g. a more explicit extra-dimensional set shift^34^. Given that the broader construct of control depends on a distributed fronto-striatal network^18,52^, there are likely multiple access points into that network. Broader explorations, including in animal models^49^, may clarify which stimulation targets are most useful in which situations.

Based on two participants reporting subjective well-being during our experiments, it seems possible that improved control might be directly reflected in standard mood/anxiety scales, but this would be a very noisy metric. Objective measures of cognitive control would also be important to ensure that the first use of such a technology rests on strong ethical grounds. Patients have a strong interest in closed-loop neurotechnologies^53^, but there are broad societal concerns about altering personalities or authentic selves through such technologies^54,55^. Given that we could improve task performance in patients with no overt cognitive impairment, these results also raise concern for attempts at cognitive enhancement in healthy humans. Given the numerous concerns surrounding potential enhancement^56^, there should be careful ethical consultation before this approach is used outside of a rigorous and restricted research setting.

In conclusion, we have developed methods for real-time monitoring of human cognitive control task performance, detection of lapses, and closed-loop remediation of those lapses. We produced potentially useful cognitive and emotional effects with far less energy than conventional deep brain stimulation. The same framework could also be applied to other cognitive/emotional problems, e.g. monitoring and enhancement of learning^57,58^ or emotion dysregulation^59,60^. Although substantial challenges remain before these results can be directly applied in the clinic, they could be the basis of a novel and highly specific approach for intervention in human neuro-psychiatric disease. In theory, this could eventually lead to application in a wider range of disorders than existing neurostimulation therapies.

## Data/Code Availability

On publication, all analysis code will be available via our laboratory GitHub. Pre-processed and anonymized neural data will be available through OpenNeuro. The closed-loop neurostimulation system has been released as open-source code with a corresponding preprint^45^, and the neural decoding and state-space modeling engines have similarly been released for open download (https://github.com/TRANSFORM-DBS/Encoder-Decoder-Paper and https://github.com/Eden-Kramer-Lab/COMPASS).

## Acknowledgements

We gratefully acknowledge technical assistance with data collection from Afsana Afzal, Gavin Belok, Kara Farnes, Julia Felicione, Rachel Franklin, Anna Gilmour, Aishwarya Gosai, Mark Moran, Madeleine Robertson, Christopher Salthouse, Deborah Vallejo-Lopez, and Samuel Zorowitz. We also thank the research participants, without whose generous help none of this would have been possible. This work was supported by grants from the Defense Advanced Research Projects Agency (DARPA) under Cooperative Agreement Number W911NF-14-2-0045 issued by the Army Research Organization (ARO) contracting office in support of DARPA’s SUBNETS Program, the National Institutes of Health (UH3NS100548, R01MH111917, R01MH086400, R01DA026297, R01EY017658, K24NS088568), Ellison Foundation, Tiny Blue Dot Foundation, MGH Executive Council on Research, OneMind Institute, and the MnDRIVE and Medical Discovery Team-Addictions initiatives at the University of Minnesota. The views, opinions, and findings expressed are those of the authors. They should not be interpreted as representing the official views or policies of the Department of Defense, Department of Health & Human Services, any other branch of the U.S. Government, or any other funding entity.

## Author Contributions

ASW, DDD, ENE, and SSC designed the study. IB, AY, BC, RZ, and UTE designed key software and tools required for data collection. KKE and TD selected the psychometric scales administered to participants and provided unpublished data related to norming of those questionnaires. ENE and GRC performed all surgical procedures. ASW, IB, BC, RZ, ACP, SSC, and DSW collected data with participants during acute seizure monitoring. ASW, IB, AY, ACP, and NP analyzed data. IB and ASW wrote the paper with substantial inputs from AY, RZ, ACP, and SSC. All authors had opportunities for critical input into and revision of the submitted manuscript, and approved its submission.

## Methods and Materials

### 1. Experimental Design

Twenty-one participants (age range: 19-57, mean age: 35, female: 12/21, left handed: 5/21) with long-standing pharmaco-resistant complex partial seizures were voluntarily enrolled after fully informed consent according to NIH and Army HRPO guidelines. Consent was obtained by a member of the study staff who was not the participant’s primary clinician. Study procedures were conducted while participants underwent inpatient intracranial monitoring for seizure localization at Massachusetts General Hospital or Brigham & Women’s Hospital. The electrode implants were solely made on clinical grounds and not tailored for research purposes. Informed consent and all other ethical aspects of the study were approved and monitored by the Partners Institutional Review Board. Participants were informed that their involvement in the study would not alter their clinical treatment in any way, and that they could withdraw at any time without jeopardizing their clinical care. This was an exhaustive sample of all participants who were available, consented to research, and had electrodes in brain regions of interest. There was no pre-planned power calculation. Data collection ceased at the end of a continuous 4-year period once study funding lapsed. Detailed clinical information on participants is in Table S1.

The core hypothesis, that internal capsule stimulation would enhance cognitive control (shorten response times in a cognitive control task without altering error rates) was pre-specified based on our prior work^17^. The analyses of behavioral and electrophysiological data, including the state-space and neural decoding modeling described below, were similarly pre-planned. The analyses described up through main text Figure 2 were specified to replicate the prior study and demonstrate that its effects were robust to changes in study population and stimulation details. Participants were part of a larger multi-modal study of the neural basis of mental illness^12^. They performed multiple other cognitive tasks both before and during their hospital stay and completed self-report scales related to emotional and cognitive difficulties. The list of tasks and scales and our strategies for comparing study participants against a normative database are described in ^12^. All are well-validated assessments that have previously been used in large-scale studies. The specific comparison of these scales against participants’ experience of subjective improvement was not pre-planned, as we did not know in advance that participants would report these effects.

Since we did not have a pre-specified stopping rule, we performed a *post hoc* sensitivity analysis in G*Power 3.1^61^ to verify that we were adequately powered for the pre-specified hypothesis that capsular stimulation would improve cognitive control. The regression coefficients reported in Tables S2 and interpreted as significant in the text correspond to a partial R^2^ of 0.0045 for the RT model and 0.01 for the theta-power model. These correspond to effect sizes f^2^ of 0.0045 and 0.01, respectively. With the sample counts/degrees of freedom as in those tables, we have over 87% power for the detected effect sizes.

### 2. Behavior Paradigm – Multi Source Interference Task (MSIT)

Participants performed the Multi-Source Interference Task (MSIT) ^62^ with simultaneous recordings of behavior and local field potentials (LFPs) from both cortical and subcortical brain structures. MSIT is a cognitive control task known to induce statistically robust subject-level effects, at both the behavioral and neural level^17,62,63^. These relatively large effect sizes amplified our ability to detect stimulation-induced differences, by increasing task-related behavioral and neural signatures. MSIT trials consisted of three numbers between 0-3, two of which had the same value (Figure 1A). Participants had to identify, via button press, the identity of the number that was unique, not its position. Each trial contained one of two levels of cognitive interference/conflict. Low conflict or congruent (C) trials had the unique number in a position corresponding to its keyboard button, and flanking stimuli were always ‘0’, which is never a valid response. High conflict, or incongruent trials (I), had the position of the unique number different from the keyboard position, requiring execution of a non-intuitive visuo-motor mapping (Simon effect). On high conflict trials, the non-unique numbers were valid responses (flanker effect). To reduce the formation of response sets, we pseudo-randomized the trial sequence such that more than two trials in a row never shared the same interference level or correct response finger. This forces frequent strategy shifts and increases attention demands, which in turn increase the need to engage/deploy cognitive control. Each participant performed 1-3 sessions of MSIT. Each session consisted of multiple blocks of 32 or 64 trials, with brief rest periods in between blocks. These periods ranged from 0.9 to 57.6 minutes; the median break time was 5.25 minutes and 89.6% were under 10 minutes. During blocks, participants were instructed to keep their first through third fingers of their right hand on the response keys corresponding to the numbers 1-3. They were instructed to be as fast and as accurate as possible. Stimuli were presented for 1.75 seconds, with an inter-trial interval randomly jittered within 2-4 seconds. Stimuli were presented on a computer screen with either Presentation software (Neurobehavioral Systems) or Psychophysics Toolbox^64^.

### 3. Electrophysiologic Recording

We recorded local field potentials (LFP) from a montage of 8-18 bilaterally implanted depth electrodes (Figure 1C, left, and Figure S1). The decision to implant electrodes and the number, types, and location of the implantations were all determined on clinical grounds by a team of caregivers independent of this study. Depth electrodes (Ad-tech Medical, Racine, WI, USA, or PMT, Chanhassen, MN, USA) had diameters of 0.8– 1.0 mm and consisted of 8-16 platinum/iridium-contacts, each 1-2.4 mm long. Electrodes were localized by using a volumetric image coregistration procedure. Using Freesurfer scripts (http://surfer.nmr.mgh.harvard.edu), the preoperative T1-weighted MRI (showing the brain anatomy) was aligned with a postoperative CT (showing electrode locations). Electrode coordinates were manually determined from the CT^65^. The electrodes were then mapped to standard cortical parcels/regions using an automatic, probabilistic labeling algorithm^66^. Intracranial recordings were made using a recording system with a sampling rate of 2 kHz (Neural Signal Processor, Blackrock Microsystems, US). At the time of acquisition, depth recordings were referenced to an EEG electrode placed on skin (either cervical vertebra 2 or Cz).

### 4. Open-Loop Capsule Stimulation

We delivered electrical stimulation to either the dorsal or ventral internal capsule and surrounding striatal nuclei (Figure 1B, right). We stimulated only one site in each block. We compared dorsal and ventral stimulation because this portion of the internal capsule has a well-described dorso-ventral topography^29–31^. More ventral capsule fibers tend to originate in more ventral aspects of PFC, particularly orbitofrontal cortex (OFC). More dorsal fibers, in contrast, tend to originate in dorsolateral and dorsomedial PFC and dorsal cingulate. The latter structures are strongly implicated in cognitive control, particularly during conflict tasks^18–20,37,52^. Thus, we hypothesized that dorsal stimulation would be more effective than ventral in this specific task. We did not have a pre-specified hypothesis regarding left vs. right stimulation, but tested hemispheres separately because our past clinical studies of capsular stimulation suggested lateralized effects^67^. We did not test any other stimulation sites for their effects on MSIT performance, because we had a specific hypothesis regarding the internal capsule. Other sEEG electrodes were stimulated in separate experiments in these same participants, but those are reported elsewhere^59,68,69^ and the outcomes of those experiments did not influence the present study.

We varied the order in which we stimulated the different sites (Table S2). Stimulation experiments were performed towards the end of each participant’s stay, when he/she was back on anti-epileptic medication, to reduce the risk of evoking a seizure. Such a session always started with 1 or 2 blocks of unstimulated trials and ended with an unstimulated block (Figure 1D). In blocks with stimulation, it occurred on only 50% of the trials. The stimulated trials were chosen pseudo-randomly, with no more than 3 consecutive trials being stimulated. All participants received the same pseudo-random order of stimulated/unstimulated trials. The stimulation was a 600 ms long train of symmetric biphasic (charge balanced) 2-4 mA, 90 µs square pulses at a frequency of 130 Hz. Stimulation was delivered through a neighboring pair of contacts on a single depth electrode (bipolar), with the cathodal (negative) pulse given first on the more ventral contact. Stimulation was delivered by a Cerestim 96 (Blackrock Instruments), with parameters set manually by the experimenter and stimulation triggered by a separate PC that was either delivering or monitoring task/behavioral stimuli.

The stimulation frequency was chosen based on a previous study^17^; it is also the frequency most commonly used in clinical DBS for psychiatric disorders. In contrast to that study, here we were able to harmonize stimulation parameters between participants, because stimulation was not directly linked to medical treatment. All stimulation was delivered at the image onset to influence a decision-making process that begins with that onset. In all participants, before task-linked stimulation, we first tested stimulation at 1, 2 and 4 mA, for 1 second, repeated 5 times with 5-10 seconds between each 130 Hz pulse train. We informed participants that stimulation was active and repeatedly asked them to describe any acute perceptions. We verified a lack of epileptiform after-discharges and ensured that the participants could not detect stimulation, e.g. through unusual sensations. If participants reported any sensation (e.g., tactile experiences in limbs or head), we limited task-linked stimulation to the next lowest intensity. In a secondary analysis, the recordings during these test stimulations were used to verify that dorsal and ventral stimulation sites had different patterns of cortical activation (see Figure S3).

Before and after all stimulation experiments, including closed-loop stimulation (see below), an experienced psychiatric interviewer (ASW) asked participants to describe their current emotional state. We similarly prompted participants to describe any subjective experiences at arbitrary times throughout each stimulation experiment. All such reports were videotaped and transcribed. During the behavioral task runs, participants were fully blind to whether stimulation was active, i.e. we did not inform them that we were beginning stimulated blocks.

### 5. Behavior Data Analysis

The primary behavior readout is the reaction time (RT), as all participants were over 95% accurate. First, we analyzed the effects of stimulation and task factors at the trial level using a generalized linear mixed effect model (GLME), as in our prior work^17^:

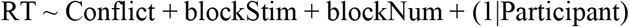

This and all other GLMEs analyzing RT data used a log-normal distribution and identity link function. Fixed effects in the GLME were Conflict (a binary variable coding the trial type as being low (0) or high (1) conflict), stimulation site (blockStim), and block number (blockNum) to account for fatigue or practice effects. Stimulation (blockStim) was coded at the block level, i.e. whether the stimulation site in a given block was dorsal vs. ventral capsule or left vs. right, not whether stimulation was on vs. off on a given trial. Block-level coding was a more parsimonious fit to the data (Akaike Information Criterion = −449.3 for block-level coding, −359.7 for trial-level coding). Participant was a random effect. All categorical variables were automatically dummy-coded by MATLAB’s “fitglme” function. We excluded trials with missing responses and with incorrect responses.

We further tested for a possible interaction between stimulation and the trial-to-trial conflict level, by fitting an alternate model with an interaction term:

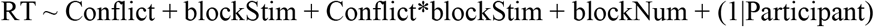

We assessed this model against our primary GLME by comparing Akaike’s Information Criterion (AIC), which decreases in models that are more parsimonious fits to the observed data.

To develop closed-loop control and neural decoding strategies, we needed to convert this block-level analysis to a trial-by-trial estimate of participants’ RT. Because the task rapidly switches back and forth between trial types, however, RT on any given trial is influenced by changes in conflict/interference in addition to (putative) stimulation effects and random variability. We sought to separately measure these processes and their change in response to capsular stimulation. To achieve this, we applied a state space or latent variable modeling framework^70,71^. In this type of model, the variables of interest (here, the “true” baseline RT and conflict-induced slowing, without influence from stochastic RT variations) are not directly observable. They are assumed to influence an observable variable (the actually observed RT) through a functional scaling, with additive noise. A further advantage is that this class of models is Markovian – the estimate of the unobserved variables (“states”) on any given trial depends only on the currently observed RT and the estimate of the states on the prior trial. This allows efficient, trial-by-trial computation that is well suited to real time monitoring and control. For exactly this reason, a specific class of state-space model, the Kalman filter, has long been used for neural decoding in motor brain-computer interface applications^35,36,72^.

Here, we used the COMPASS toolbox^71^ for MATLAB to fit a model that extracts a trial-level estimate of the RT independent of conflict effects. This model takes the form:

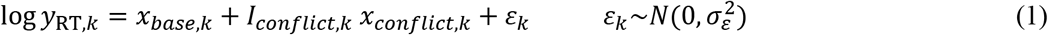

where *y*_RT,*k*_ is the RT on trial *k*, and the *x*_*k*_ are latent, unobserved variables that we have previously termed “cognitive states”. The observation noise, *ε*_*k*_ would capture other non-structured processes that influence the trial-to-trial RT. Note that this model follows the same distribution/link assumptions as the static GLME above, namely a log-normal distribution. One of the differences between this modeling approach and the classic Kalman filter is that RT is markedly non-Gaussian. It can only have positive values, and is very skewed – RT distributions across a range of cognitive tasks have long right tails^71,73,74^. The log-normal assumption corrects for this.

The latent variables were modelled as:

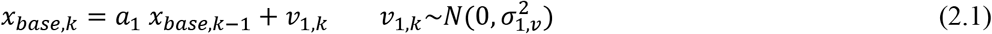

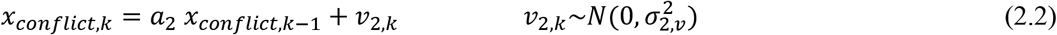

where, *a*_1_ and *a*_2_ define the decay of the state variables over time. *v*_1,*k*_ and *v*_2,*k*_ are mutually independent white noise processes with zero mean and variance 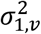 and 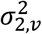, respectively. That is, we assumed that these two processes can vary entirely independently of one another (even though stimulation may influence both). Because both the state values on each trial and the parameters linking those values to the RT are unknown, this model has no closed-form solution. COMPASS estimates model parameters through an iterative expectation-maximization algorithm^71^. We verified convergence of this algorithm on all participants by inspection of the model likelihood plot. We also tested models that included both accuracy and RT as observable outputs; these yielded no significant improvement over RT-only models (see main text and Figure S8).

*x*_*base,k*_ represents the expected RT in the absence of conflict or other external influencing factors, whereas *x*_*conflict,k*_ represents the expected effect of conflict on the RT. *I*_*conflict,k*_ is an indicator variable, such that *x*_*conflict,k*_ only affects the expected RT on high-conflict trials. *x*_*base*_ can be thought of as encoding more general, overarching aspects of cognitive control, such as effortful attentional focus on task stimuli, maintenance of goals in working memory, and preparation to inhibit a prepotent response on incongruent/high-conflict trials. *x*_*conflict*_, in that framework, represents the cognitive load of actually deploying the response inhibition in response to conflict. This is sometimes framed as a “reactive” model of cognitive control. We derived this two-variable model from both theory and prior work. We previously observed^17^ (and confirmed in this work, see above) that capsular stimulation affects both high- and low-conflict trials. That is, it mainly alters the expected RT *x*_*base*_. We considered, however, that reactive control in response to conflict might also be affected, and wanted to specifically test for that potential effect. Further, in prior work, we found that this two-state model was necessary to accurately model RT during this same task^70^. We verified that the two-state model was an appropriate fit to our data by comparing the RT residuals to those expected from a white-noise process. Unmodeled variance (e.g., a third cognitive process not captured in our model) would lead to structured residuals that violate white-noise expectations.

The state-space model assumes that cognitive states are slowly varying, i.e. they show a strong autocorrelation. (This was true in practice; the estimated values of *a*_1_ and *a*_2_ were close to 1 for all participants.) We thus cannot use the GLME to analyze stimulation-induced change in these latent variables (*x*_*base*_, *x*_*conflict*_) because they strongly violate the GLME’s assumption that individual datapoints are independent. We instead used non-parametric permutation testing, which is well-established as a method for inferential statistics on autocorrelated time-series^75^. The stimulation labels of individual blocks were shuffled 1,000 times, with the shuffling nested within individual participants. This created a distribution of cognitive state values under the assumption of no difference between stimulation sites (or between stimulated and non-stimulated trials). From that distribution, we inferred the p-value of the actual state values under stimulation. For both the raw RT GLME and the cognitive state permutation tests, we compared up to 4 stimulation sites in each participant to baseline (no stimulation, NS1). Within each analysis, we corrected the p-values for these multiple comparisons using a false discovery rate (FDR) step-down procedure via MATLAB’s “fdr” function.

### 6. Closed Loop Stimulation

In three participants, we used that same state space model to implement closed-loop control. First, for each participant, we estimated model parameters by running the expectation-maximization fitting algorithm on 1-3 days of prior MSIT performance without brain stimulation. We used all available data for each participant to fit these models. These parameters were then provided to a real-time engine (also based on COMPASS, specifically the real-time decoding function, compass_filtering) that estimated *x*_*base*_ and *x*_*conflict*_ on each trial. This specifically leverages the Markovian nature of our model, in that estimating these states on each new trial is extremely fast once the model parameters are initially estimated.

We attempted to control *x*_*base*_, which we considered to track the overall difficulty of sustaining attention and exerting cognitive control during conflict (more difficulty leading to longer RTs). In that framework, cognitive control enhancement would be reflected in a decrease in *x*_*base*_. To achieve this, if the estimate on trial *k* was above a manually selected threshold, the system delivered electrical stimulation at the time of image/stimulus presentation on trial *k+1*. The threshold and stimulation effects were visible to the experimenter, but not to the participant, through a custom GUI running on a separate laptop. We set the threshold to attempt a decrease in *x*_*base*_ from its unstimulated value. The supervising experimenter (ASW) observed the participant’s best performance from real-time state estimation during the initial non-stimulation block (64-128 NS1 trials). This process estimated both the mean (maximum likelihood value) of *x*_*base*_, plus a confidence bound. We then set the threshold/target for *x*_*base*_ to be below that lower confidence bound, i.e. to be outside the range of possible values that were consistent with the baseline/unstimulated performance. There was no quantitative rule for the distance between the estimated state bound and the control target; it was selected *ad hoc* to be outside our subjective estimate of overall process variability. Stimulation parameters and hardware were identical to the setup for open-loop stimulation. Because of limited experimental time, we could do either open-loop or closed loop stimulation experiments with a particular participant, but not both. We attempted closed-loop control using the stimulation site that had the largest behavioral effect during open-loop experiments in the 6 previous participants. When that site was unavailable (not implanted in a given participant), we used the next best choice that was available.

For analysis of the closed loop stimulation results, we re-ran the complete state-space estimation offline over the whole dataset, rather than using the less-accurate state values estimated in real time. A key difference is that the offline estimation contains a forward (filtering) and backward (smoothing) pass, allowing future data to influence each trial’s estimate non-causally. By considering more information, this offline estimate more accurately reflects the “true” cognitive process and its change in response to stimulation. To directly compare closed-loop and open-loop stimulation, we normalized the state values between these two runs such that that the unstimulated blocks in both paradigms had a mean value of 1. That is, both open-loop and closed-loop results were expressed as change vs. the unstimulated condition on the same day.

### 7. Neural Data Analysis - Preprocessing

Local field potentials (LFP) were analysed using custom analysis code in MATLAB (Mathworks) based on FieldTrip (http://www.fieldtriptoolbox.org/). To reduce the influence of volume conduction^76^, LFPs were bipolar re-referenced by subtracting those recorded at consecutive electrode contacts on the same electrode shank. LFP was recorded from electrode pairs spanning 16 brain regions: prefrontal, cingulate, orbitofrontal, temporal, and insular cortices, amygdala, hippocampus, nucleus accumbens, and caudate (Figure S1). All LFP data were decimated to 1000 Hz and de-meaned relative to the entire recording. 60 Hz line noise and its harmonics up to 200 Hz were removed by estimating noise signals through narrow bandpass filtering, then subtracting those filtered signals from the original raw signal. We removed pathological channels with interictal epileptiform discharges (IEDs). We detected such channels with a previously-reported algorithm that adaptively models distributions of signal envelopes to discriminate IEDs from normal LFP^77^. We then used a Morlet wavelet decomposition to estimate power in 6 frequency bands (4-8, 8-15, 15-30, 30-55, 65-110, and 135-200 Hz). This decomposition used a 1 Hz frequency resolution and 10 ms timestep, with the default parameters of 7 cycles per wavelet and a Morlet/Gaussian width of 3 standard deviation. We fractionated the high gamma (65-200 Hz) band into lower and upper bands to bypass the stimulation frequency at 130 Hz and a 60Hz harmonic at 120 Hz. All time-frequency decomposition was performed after individual trials/epochs were cut from the continuous recording, so that artifacts on stimulation trials could not influence power calculations on non-stimulation trials. Each trial was cut with a buffer of 2 seconds on each side to mitigate edge effects in the wavelet transform; data from these buffers were discarded before analysis.

### 8. Neural Data Analysis – Mid-Frontal Theta Power

Exercise of cognitive control is associated with higher theta (4-8 Hz) power in a fronto-cingulate network^17,19,27,28,78^. We have specifically reported that stimulation in the internal capsule increases that task-evoked theta^17^. As an initial manipulation check, we sought to replicate these prior results. We analyzed an epoch of 0.1-1.4 seconds after image onset, which covered the decision-making period up to the median RT. We focused this analysis on non-phase-locked oscillations, which are more strongly correlated with cognitive conflict effects than are their phase-locked counterparts^24,79^. From the target epoch, we subtracted the time-domain evoked response (ERP), which contains all phase-locked activity. We calculated this ERP separately for high- and low-conflict trials, and subtracted the appropriate ERP from each trial’s time-domain data. We then transformed the time-domain to a time-frequency representation as above. We averaged power in our analysis epoch within the theta band. For visualization, we normalized this power as a log ratio relative to a baseline period of 0.5 seconds preceding image onset. For analysis, this log transformation is built into the GLM (see below).

To verify that higher conflict evoked higher frontal theta, we analyzed the blocks without stimulation. This avoids confounding effects of stimulation and conflict. For each participant, we pre-selected pre-frontal cortical (PFC) channels that had a significant increase over baseline in task-evoked theta (t-test with threshold of p<0.05 uncorrected). For this initial pre-screening step, to avoid a circular analysis, we did not split trials into high/low conflict. Rather, we identified channels that showed a theta-band response in general to performing MSIT. In this reduced set of channels, we then divided the trials into low and high conflict, then computed the non-phase-locked theta power, as noted above. We combined all pre-selected channels in each PFC region, and for each region we fit the GLME:

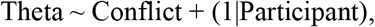

where Conflict is a binary variable coding the trial type as being low (0) or high (1) conflict. This and all other GLMEs analyzing LFP power data used a log-normal distribution and identity link function. We false discovery rate (FDR) corrected the resulting p-values for testing of multiple PFC regions.

We then tested whether open loop capsular stimulation caused a significant increase in theta in the unstimulated trials within a stimulation block (NS2) compared to those in the unstimulated blocks (NS1; see Figure 1C-D). To accurately assess stimulation effects, we discarded stimulation trials that we presumed to be substantially contaminated by artifact. We then compared two types of non-stimulated trials (Figure 1D). NS1 trials were from blocks in which no brain stimulation was given on any trial. NS2 trials were from blocks with stimulation, but were the pseudo randomly-selected 50% of trials that did not receive stimulation. These NS2 trials were artifact-free, but still showed the behavioral effect of stimulation, and thus should also show physiologic changes related to that behavior change. We therefore tested whether the normalized theta power in NS2 trials was significantly greater than that in NS1, again using a GLME:

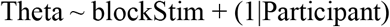

For this model, we chose one PFC channel for each participant that had the highest theta during NS2 trials (regardless of conflict level or stimulation site, again to avoid circular analysis). P-values were again FDR corrected to control for testing of multiple stimulation sites against non-stimulation.

### 9. Neural Data Analysis - Decoding

To effectively treat psychiatric disease, closed-loop stimulation will ultimately need to be applied outside of a structured task-based paradigm, i.e. as patients go about their daily lives. That, in turn, requires the ability to detect lapses in cognitive control and conflict responses directly from brain activity. We thus developed a neural decoder for our cognitive state variables. We applied a neural encoding-decoding analysis with automatic feature selection, as in ^70^. The decoded variables were *x*_*base*_ and *x*_*conflict*_ from the model in equation (1). The neural features used for decoding were the LFP power, in the above-mentioned frequency bands, averaged over a 2 second interval starting at the MSIT image onset. We broadened the analysis beyond the theta band because prior literature suggests that while theta is strongly associated with cognitive control, other frequencies also carry significant amounts of information about task performance^26^. The 2 second epoch was chosen to include both the response and post-response processing. This wider window produced smoother features with less trial to trial variance, improving decoder stability. Here, we averaged only across a 2 second time interval (200 samples) to get power features per-trial. Similar to the theta analysis, the encoder model fitting considered only NS1 and NS2 trials, to prevent the influence of stimulation artifact. We focused on LFP spectral power (rather than other potential behavioral covariates such as connectivity/coherence) because power can be efficiently computed within currently available implantable neural devices^39,80,81^. Successful decoding of task performance from power alone could pave the way for use of these closed-loop controllers in clinical settings.

Decoding analyses were performed with out-of-sample validation, using both stimulated and unstimulated MSIT datasets. For each participant’s data, 20-66% of the total trials were used to fit an encoding model (training set). These consisted of NS1 trials in unstimulated datasets and both NS1 and NS2 trials in the stimulated experimental datasets. The training trials were selected from contiguous blocks of trials that, collectively, covered the full range of the states during an experiment. The encoding model that we used is a linear model of the form Y_k_~1+βx_k_, where Y_k_ is a neural feature and x_k_ is one of the cognitive states on the k-th trial. We considered a feature to be a candidate for decoding if the modified F-statistic^70^ of the corresponding model corresponded to p < 0.01 (uncorrected). This procedure selected a set of candidate neural features that potentially independently encoded each cognitive state. The exact number of training trials for each dataset was determined as the minimum required to have a non-zero number of features selected by the encoding procedure.

Next, to reduce overfitting, we pruned the selected set of neural features. The validation set for this pruning used 21-50% of the dataset for each participant, and had 25-50% overlap with the training set. We allowed this overlap to ensure that the validation set was sufficiently large for model pruning without leaving too few trials available for the subsequent test set (see below). On this validation set, we estimated the posterior distribution of the cognitive state from the neural data, through a Bayesian filtering process similar to the one used to estimate these same latent variables from RT^70^. We calculated the root mean square error (RMSE) between the neurally decoded state and the “true” (estimated from behavior) cognitive state in this held-out test set^70^. We then sequentially dropped the feature whose removal led to the most improvement in RMSE. The final decoder was then the set of features that survived this dropping step, i.e. where dropping any further feature would increase RMSE on the validation set.

Finally, we reported the decoder’s performance on a fully held out test set, namely the 18-46% of remaining trials from each participant that had not been used for either decoder training or feature pruning. The exact number of trials used for each participant’s decoder fitting, pruning, and testing is given in Table S6. A key challenge in performing these analyses was that the latent cognitive states (*x*_*base*_ and *x*_*conflict*_) are themselves multivariate Gaussian estimates. The estimate’s value can depend on the starting point of the expectation-maximization process used to fit the state-space model. To control for this, we re-ran the behavioral estimation for each participant 1,000 times with different random seeds, producing 1,000 estimates of the underlying trajectory. (Further details are given in ^70^). We then evaluated the neural decoder’s performance based on whether its point estimate of the decoded state was within the confidence interval derived from these multiple trajectories.

We fit this encoding-decoding model separately to data from unstimulated sessions (consisting of only NS1 trials) as well as to stimulated sessions (both NS1 and NS2 trials), to determine how the encoding structure was altered by electrical stimulation. We did not include stimulated trials in this analysis, because there is a prominent stimulation artifact that makes these trials easily discriminable. In cases of stimulation-behavior correlation, behavior could be trivially decoded simply by detecting the artifact.

## Supplementary Material

**Figure S1:**
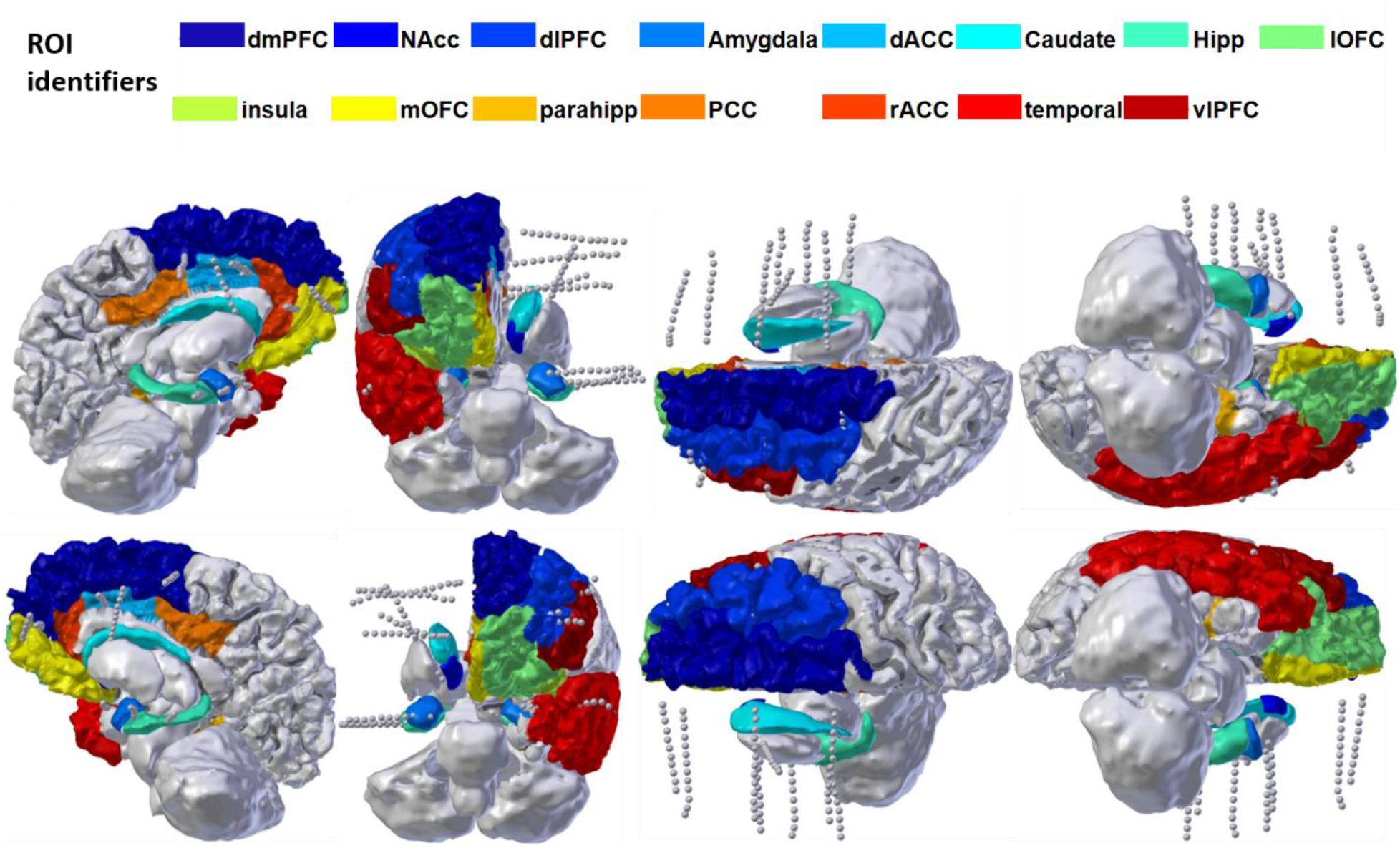
Example recording montage from a single participant, with cortical parcellation overlaid. Electrode shanks access a broad network covering multiple prefrontal structures, superficial and mesial temporal lobe, and striatum/internal capsule.

**Figure S2:**
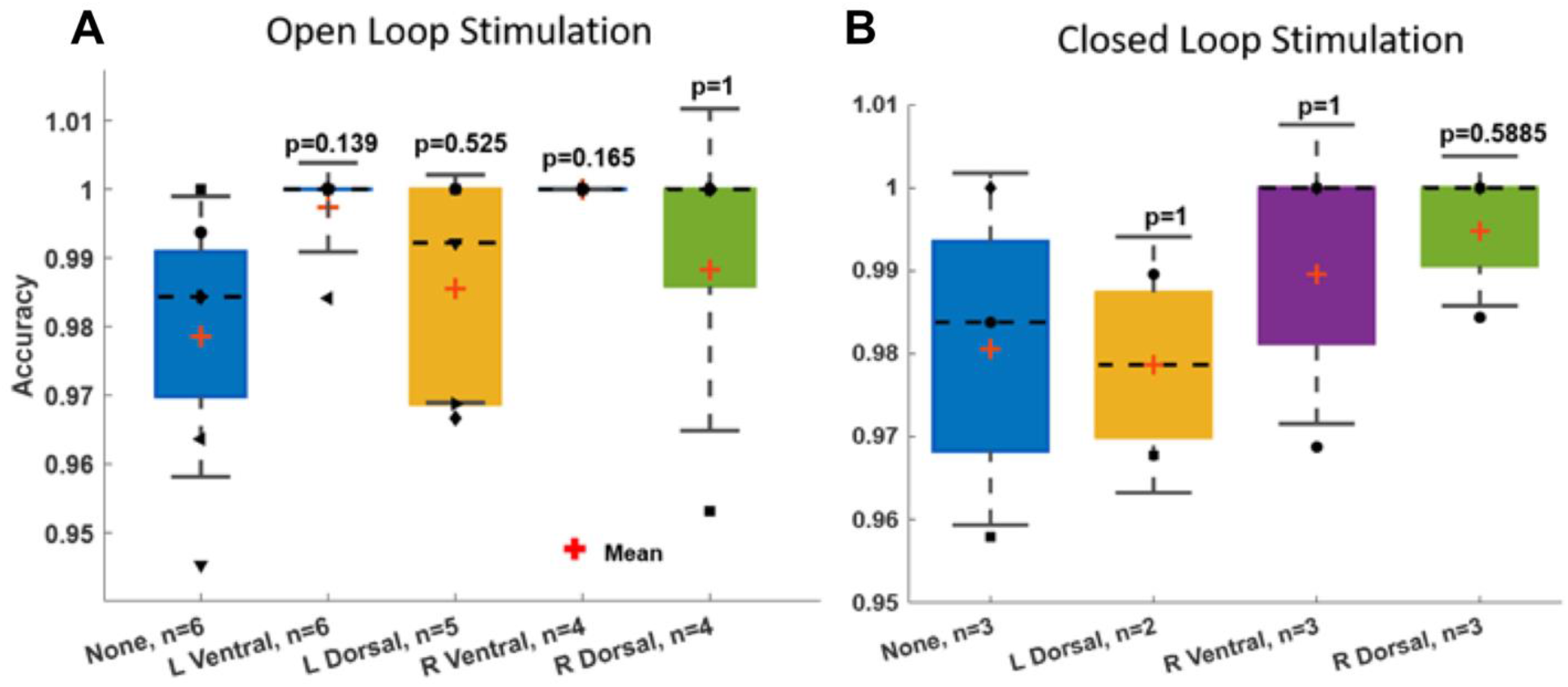
Accuracy during different stimulation experiments, for A) open-loop and B) closed-loop capsular stimulation. The p-value above each bar represents a binomial exact test of accuracy compared to the non-stimulated baseline condition, with Benjamini-Hochberg false discovery rate correction. All accuracies are above 95%, with accuracy during stimulated blocks being very slightly higher in most cases. No results exceed chance significance.

We did not have open- and closed-loop data from the same participants. To compare the CL and OL conditions, we therefore compared their accuracies across participants with a Fisher exact test for each of the three stimulation sites (L Dorsal, R Ventral, R Dorsal) that were used in both conditions.

L Dorsal: p=0.645
R Ventral: p=0.440
R Dorsal: p=0.655

These provide no evidence for a difference between OL and CL conditions.

These results do not support a change in accuracy with any stimulation type. That is, the observed decrease in reaction times is a true performance improvement, not a shift along a speed-accuracy tradeoff. We were unable to analyze accuracy in the GLME framework because the differences between stimulation sites are so small as to make the models non-identifiable in all cases.

**Figure S3:**
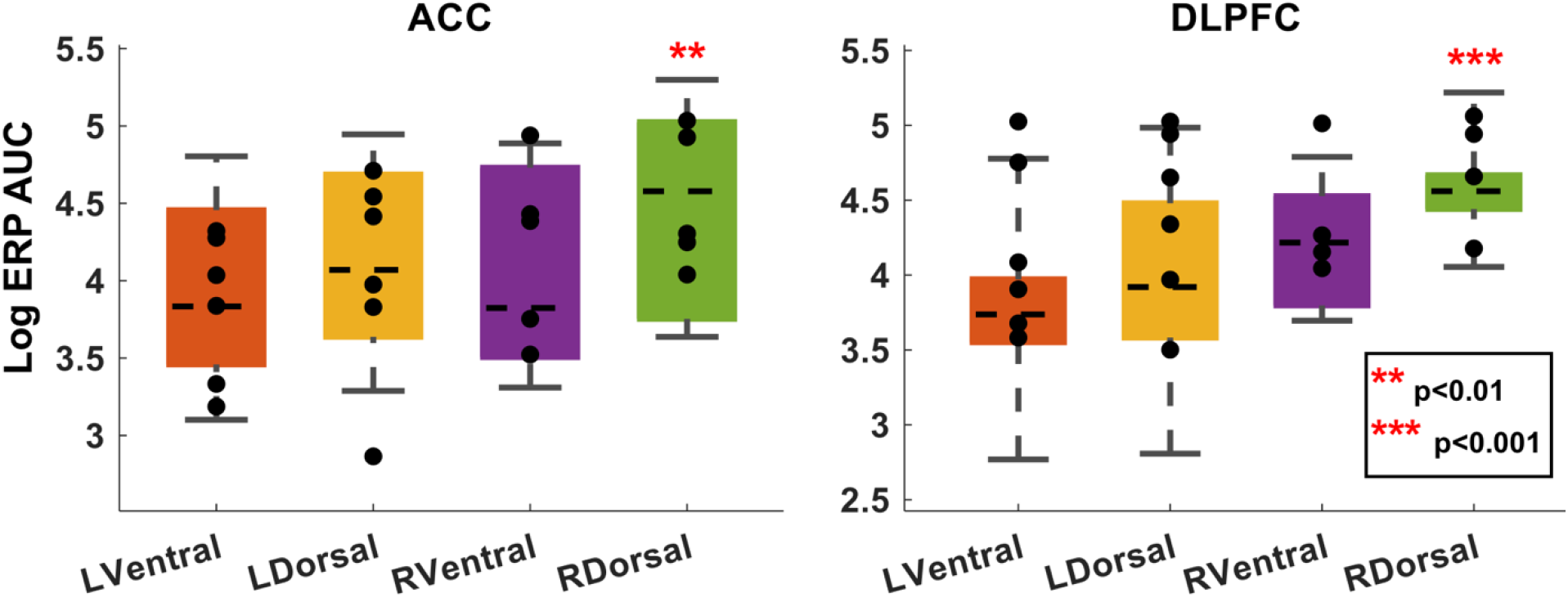
Topographic structure of the internal capsule yields differential cortical effects from stimulation at different capsular sites. Before task-linked stimulation, we performed safety/perceptibility testing, where we repeatedly stimulated each potential site with brief 130 Hz pulse trains (see Methods). Each of those trains created an evoked response potential (ERP) in various cortical regions. For each participant, we collected all sEEG channels that were localized to grey matter of DLPFC or ACC. We then quantified the post-train ERP as the sum of the area under its polyphasic curve (AUC). We limited this analysis to channels ipsilateral to the site of stimulation. Each marker represents the mean log(AUC) in one participant.

The stimulation sites that were more effective behaviorally produced the largest ERPs in these cognitive-control-associated regions, with right dorsal stimulation having the largest effects. (p-values represent t-test on the regression coefficients of a log-normal GLM, i.e. the same analysis used in main text Figure 2). In the left hemisphere, dorsal stimulation produced larger responses than ventral stimulation, but this did not reach statistical significance given the small number of trials (5 test trains per participant).

These results are consistent with the known topography of the internal capsule, where fibers that connect DLPFC and ACC to thalamus run in the dorsal-most part of the anterior limb, i.e. in close proximity to our chosen dorsal electrodes.

**Figure S4:**
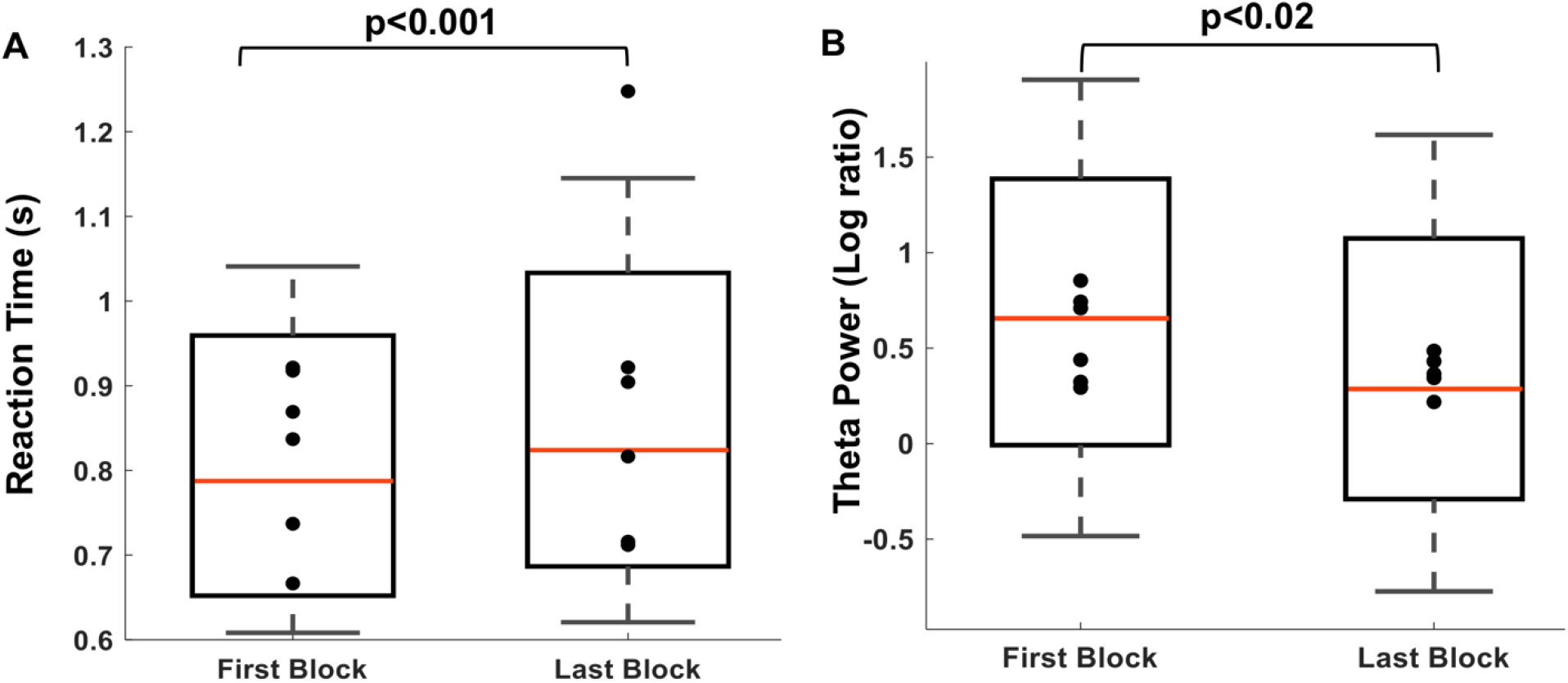
Open loop stimulation effects are not explained by practice or regression to the mean. Here we show the reaction time (A) and theta power (B) for the first and last MSIT block of each participant in the open loop experiment. Analysis windows, channel selection, and statistical testing follow main text Figure 2. All p-values are Wald tests on coefficients from a generalized linear model with Block as the primary fixed effect. Both analyzed blocks contain only NS1 trials, meaning there is no stimulation on any of the trials. In this comparison, RT significantly worsens (increases) over the course of the experiment, while PFC theta decreases. This is the precise opposite of the stimulation effects reported in Figure 2, verifying that those effects are attributable to the stimulation itself. We believe this RT increase and theta decrease represent fatigue and/or difficulty sustaining attention on task performance.

**Figure S5:**
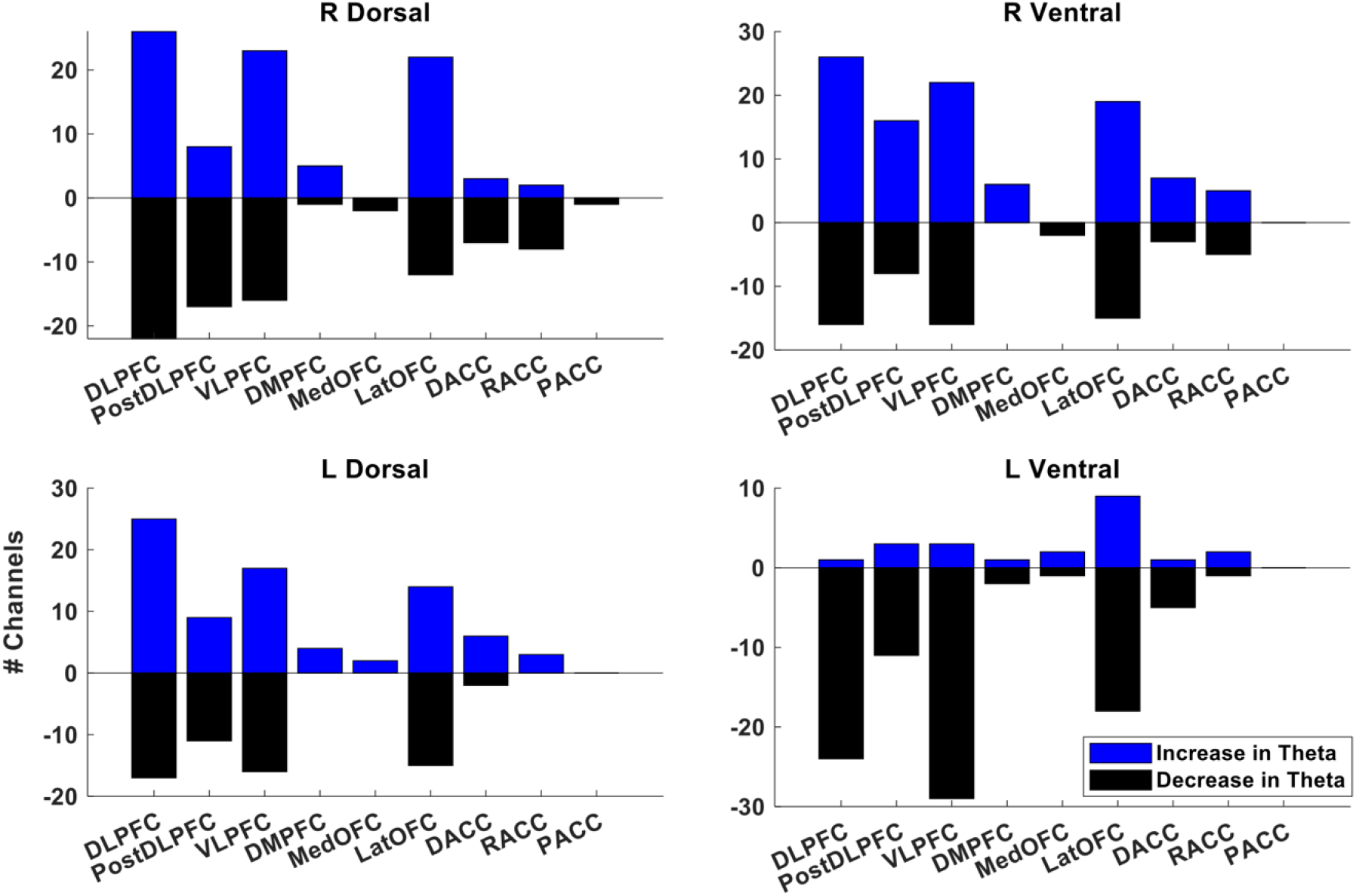
Behaviorally effective stimulation increases theta in brain regions related to cognitive control. The Y-axis plots the number of channels that are localized to the specified region, and that showed either an increase (above X-axis) or decrease (below axis) in theta power, using the same analysis window and baseline correction as in main text Figure 2. Increases and decreases are based solely on numeric comparison, not on mass univariate t-testing. To demonstrate consistency of effects across the relatively large extent of the lateral PFC, and to account for the relatively large number of channels assigned to this region, we have fractionated DLPFC into anterior and posterior sub-portions.

After stimulation at sites that were behaviorally effective (Left/Right Dorsal and Right Ventral capsule) channels with theta increases outnumbered those with decreases, specifically in regions known to be engaged by the Multi-Source Interference Task (DLPFC, VLPFC, and cingulate to a lesser degree). More importantly, after stimulation at the ineffective site (Left Ventral capsule), there were no regions where theta increases predominated. These results support a link between PFC theta augmentation and enhanced cognitive control, although they also emphasize the heterogeneity of this response and of PFC generally.

**Figure S6:**
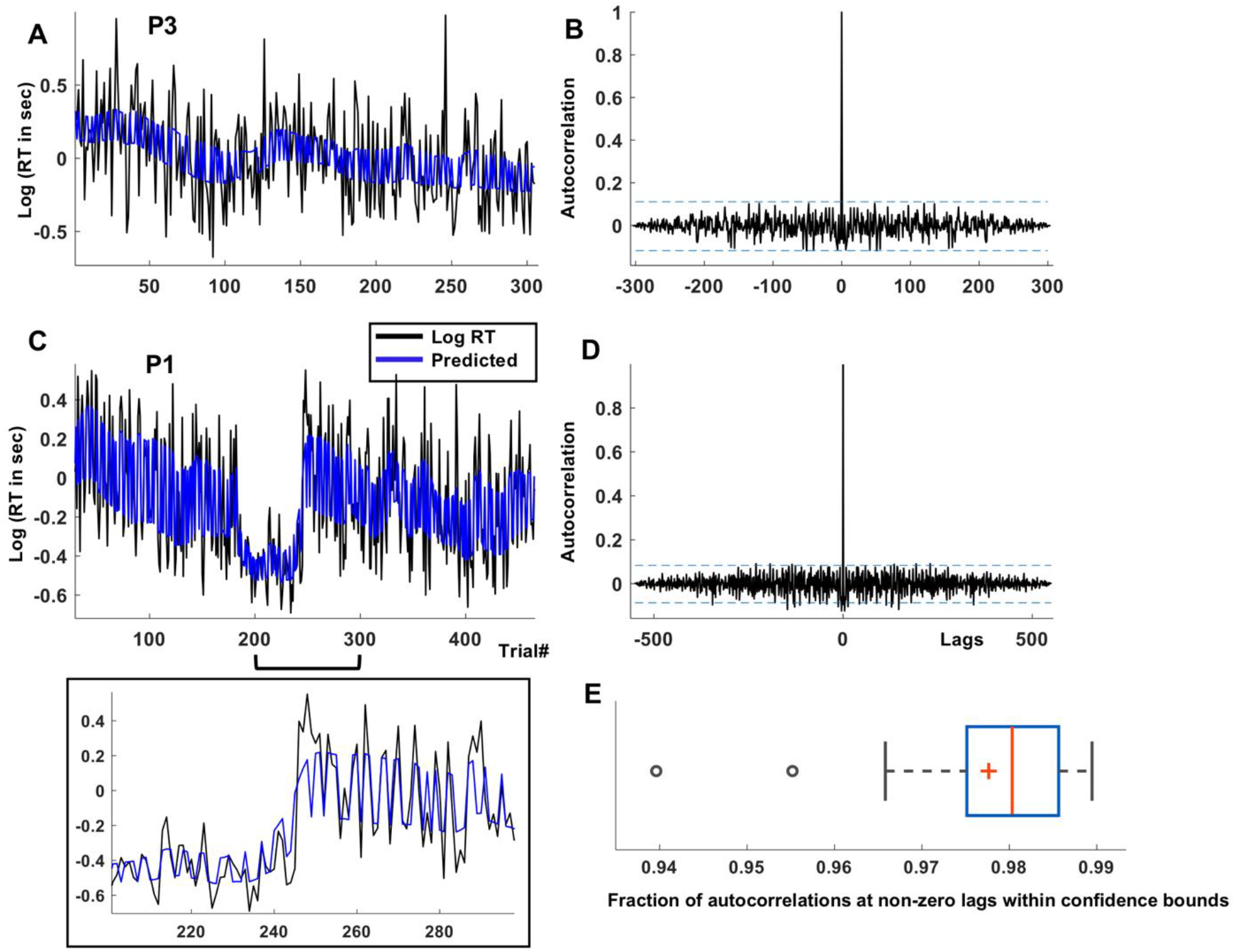
State-space model goodness of fit. A,C) Examples of actual log(RT) from one MSIT run in each of two participants (black), with model prediction superimposed (blue). The model captures both slow-scale fluctuations (primarily in *x*_*base*_) and fast, conflict-driven fluctuations (through *x*_*conflict*_) in RT. An inset below panel C provides zoomed-in detail of the two timeseries at an inflection point. B,D) autocorrelation of residual error between measured and predicted log(RT), for the same participants. The dotted blue lines indicate the confidence bound for autocorrelation at non-zero lags for a white noise process (calculated as in https://nwfsc-timeseries.github.io/atsa-labs/sec-tslab-correlation-within-and-among-time-series.html). E) Fraction of autocorrelation values at non-zero lags (as in B,D) that lay within the confidence bounds of a white noise process, for all 21 participant datasets. These are all substantially greater than 90%, indicating that the two-variable state-space model captures the primary structured sources of variance within RT.

**Figure S7:**
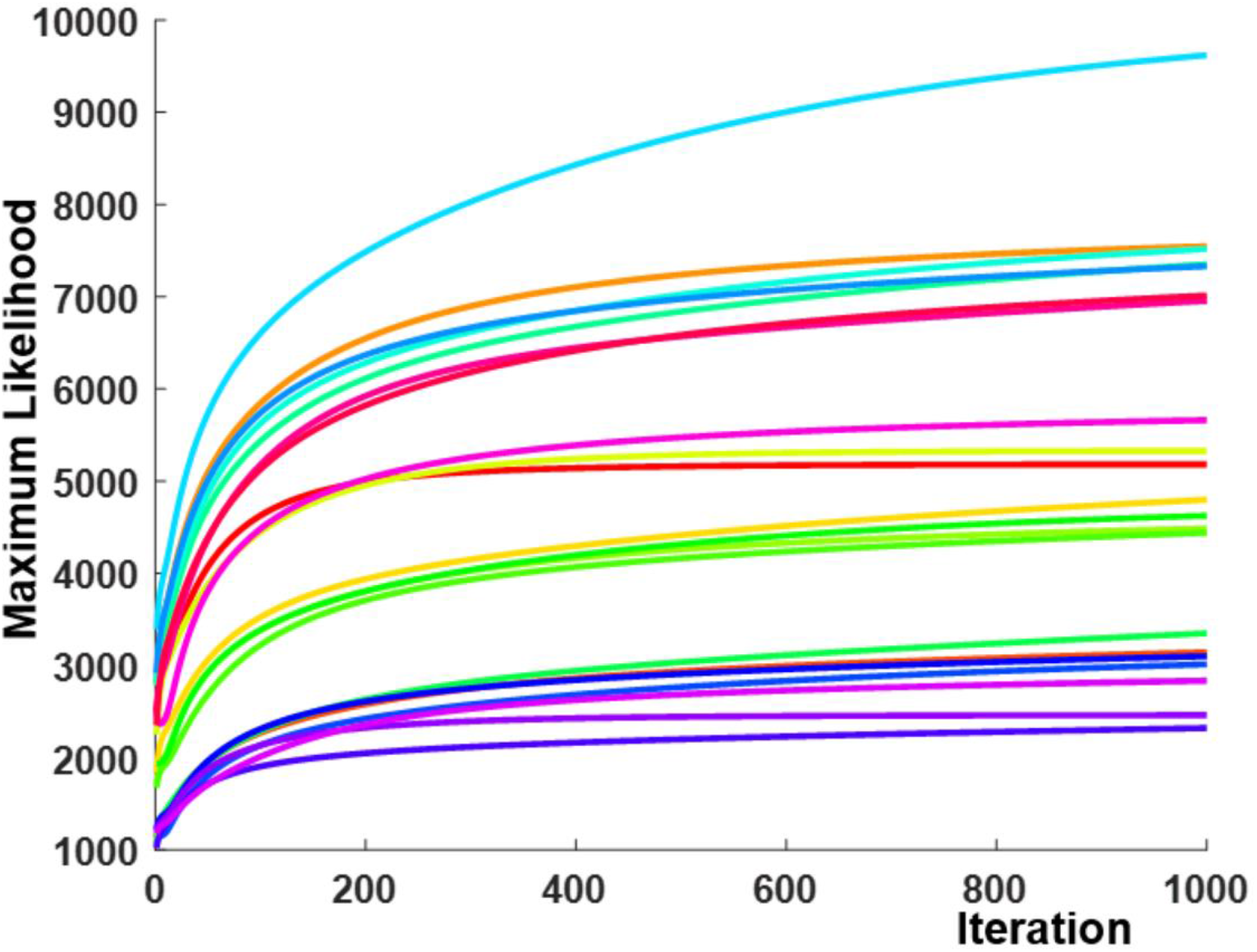
State-space modeling convergence. Each colored curve is the model likelihood, at each step of the expectation-maximization (EM) algorithm, for one participant (colors are arbitrary and do not relate to other figures). We applied a fixed termination criterion of 1000 EM iterations. By this point, all curves had reached an asymptotic maximum likelihood, and none was still in a phase of linear or supra-linear improvement.

**Figure S8:**
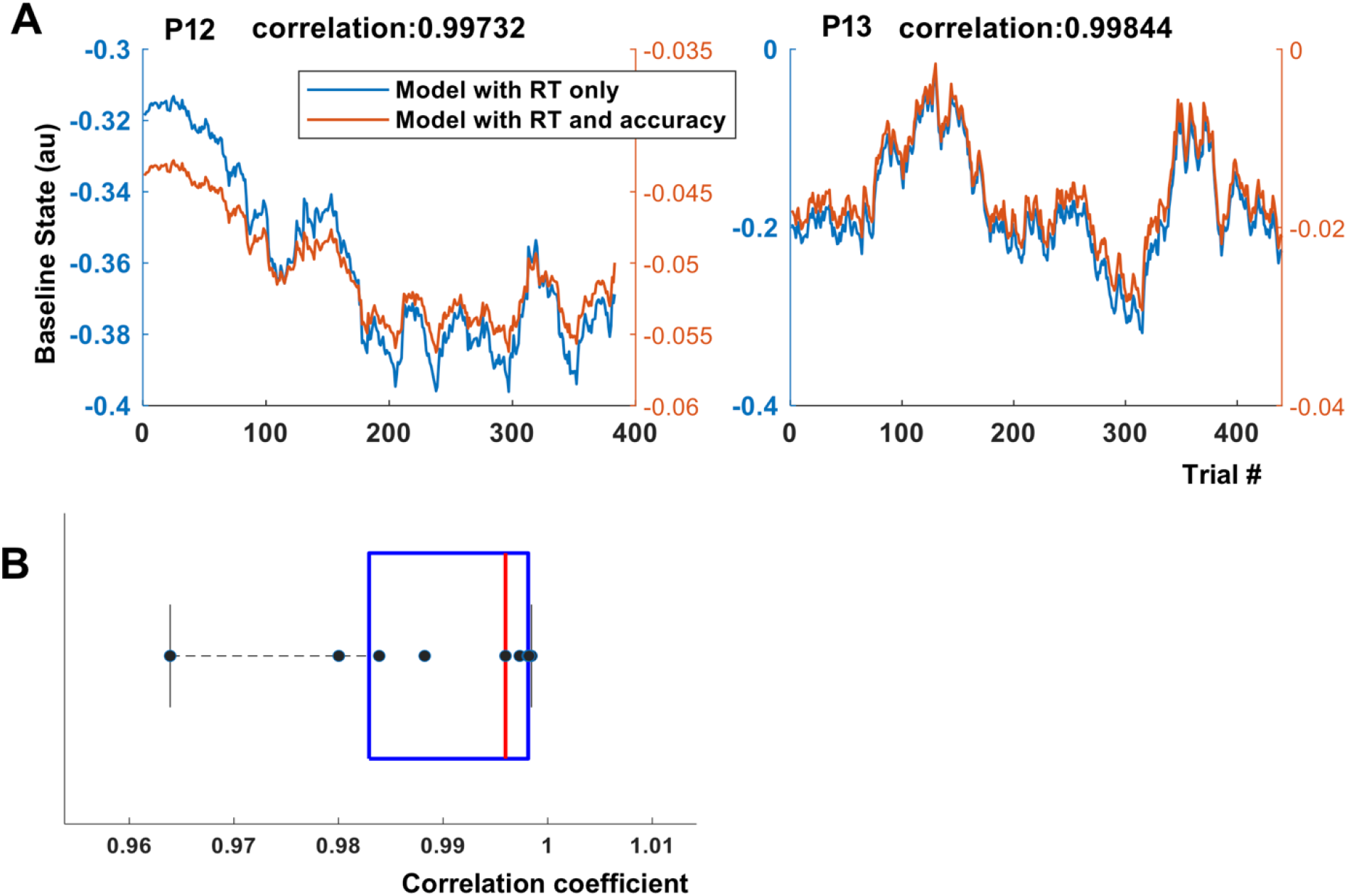
Considering accuracy in addition to RT did not improve behavioral model fitting. A), example *x*_*base*_ trajectories for models fit using only continuous RT or both RT and accuracy, in two example participants. The curves are nearly identical to within a scaling factor, as evidenced by extremely high Pearson correlation coefficients between them (above figures). B), distribution of Pearson correlations between RT-only and RT-plus-accuracy models, for all participants. All correlations are above 0.96, and the majority are above 0.99. Adding accuracy to the model would not change our inferred states or any conclusions related to stimulation effects or decoding. It would, however, add multiple free parameters to the model fitting process, offering opportunities for overfitting. This strongly argued for the more parsimonious RT-only model, which we used in all further analyses.

**Figure S9:**
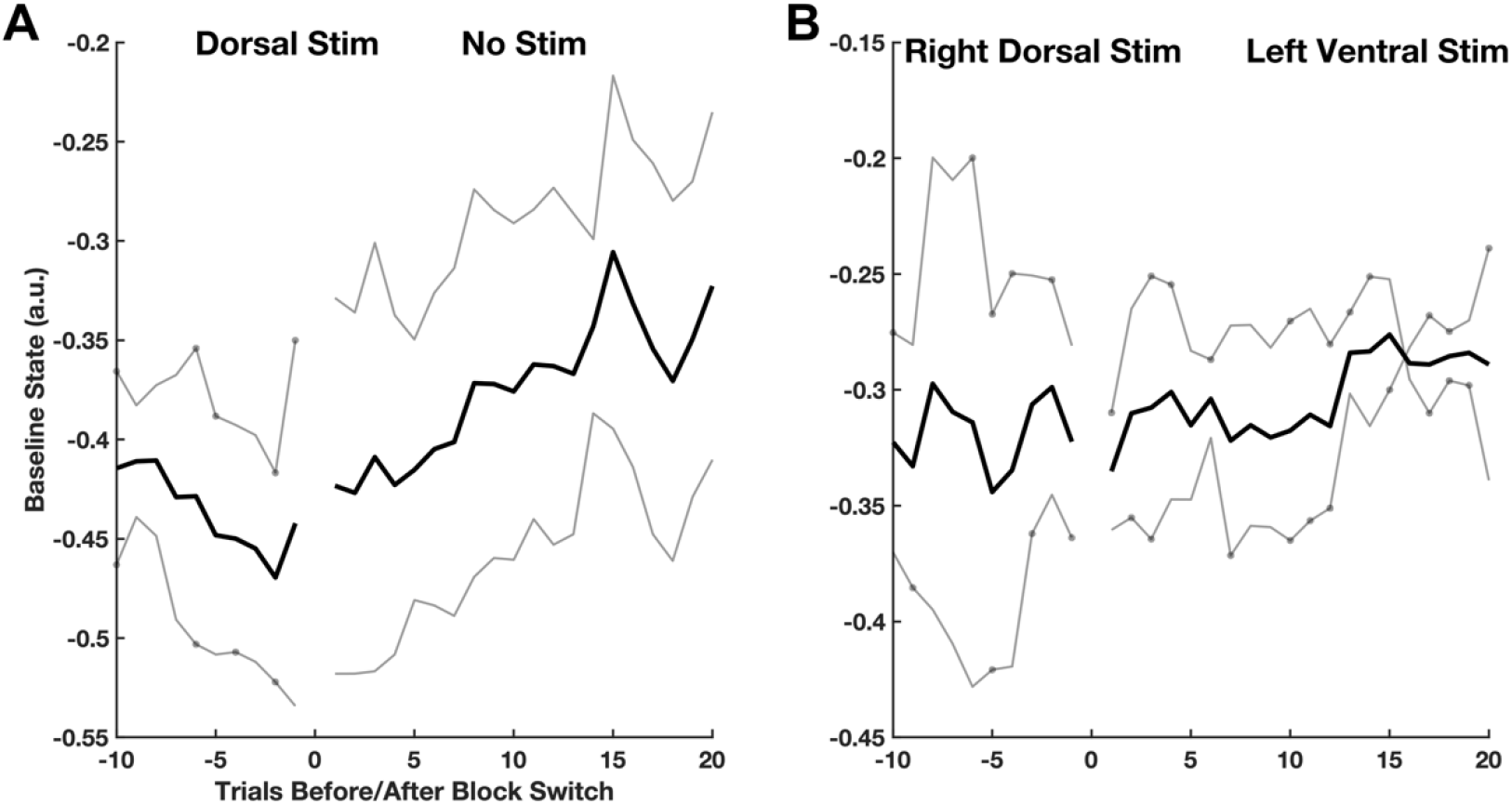
Stimulation effects had modest carry-over between blocks, but in a pattern that would not cause spurious detection of between-block differences. Each panel shows the maximum likelihood estimate of *x*_*base*_, in light grey for individual participants and dark black for the mean of all participants (n=2 in each case, based on having experienced the required type of block transition). Markers on the individual-participant curves show trials on which stimulations were delivered. A), transition from dorsal capsule stimulation (one left dorsal, one right dorsal) to the terminal non-stimulated block. *x*_*base*_ increases rapidly once stimulation ends, stabilizing at a higher value after about 10 trials. As noted in Figure S4 above, this led to a higher median RT in the final block than in the initial non-stimulated block, which we interpret as a fatigue effect. B), transition from right dorsal stimulation (our most effective condition) to left ventral stimulation (the only condition found to be ineffective in the RT analysis of main text Figure 2). For clarity, we included only participants where right dorsal stimulation was clearly different from their baseline and where this specific transition occurred (P9 and P12). Here, there appears to be some carry-over – even though left ventral stimulation was not effective in the group-level analysis, *x*_*base*_ does not rapidly increase (at best, a very subtly upward trend is visible).

We emphasize that persisting effects between blocks would not explain the behavioral or neural effects described in the main text, because these persisting/carry-over effects would tend to decrease the statistical significance and numeric size of any detected between-block differences. That is, they actively bias against the conclusions we report, suggesting that the true effects might be larger in a different experimental design.

**Figure S10:**
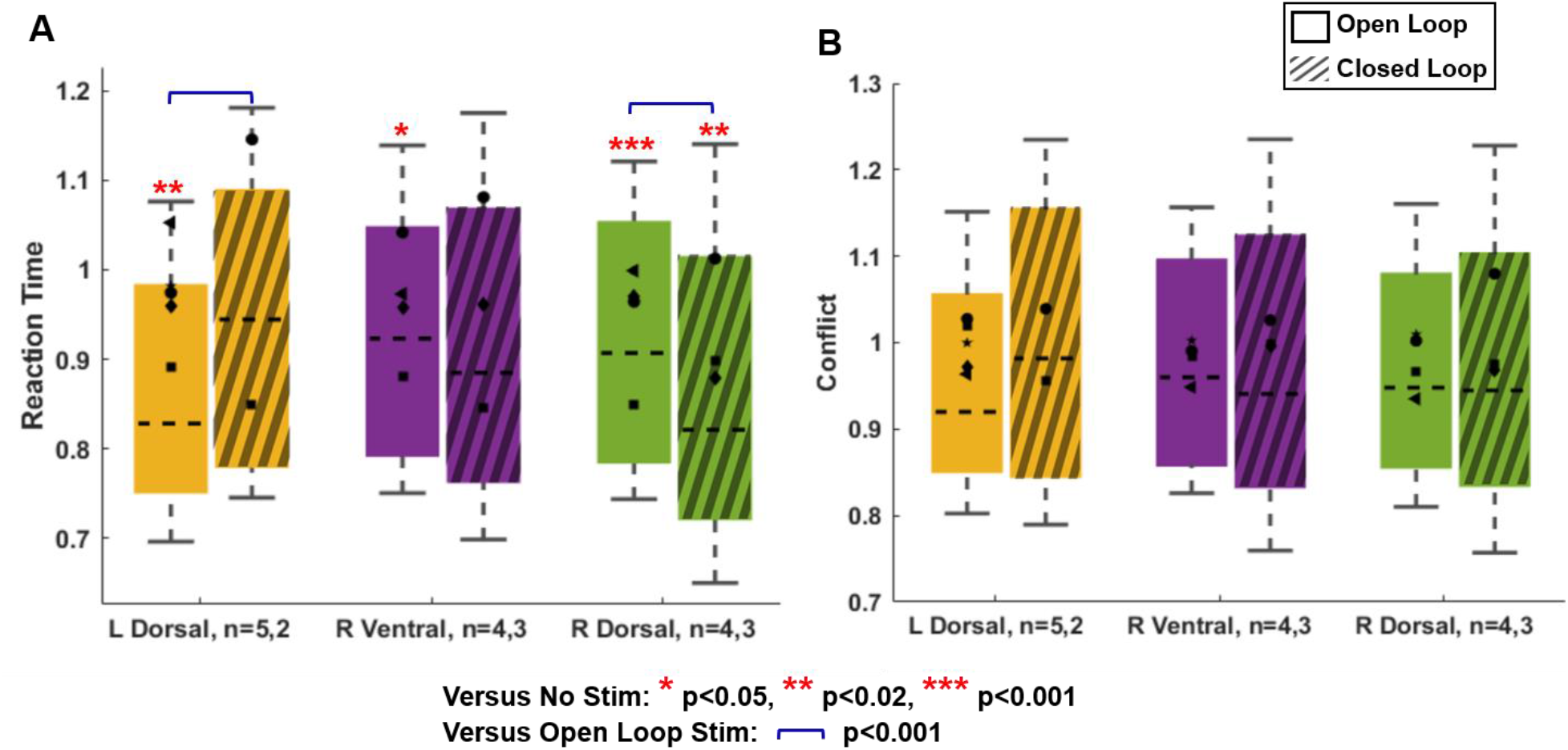
Effect of open loop and closed loop capsular stimulation on A) reaction time (RT) and B) Conflict related RT. Conflict related RT is calculated as the residual reaction time after subtracting the mean reaction time of the congruent trials in the same block, i.e. it has an expected value of 0 ms on non-conflict trials. We consider it as the closest raw/manifest data analogue of *x*_*conflict*_. We note, however, that both of these manifest RT variables include the Gaussian noise that is removed by the state-space filtering that produces *x*_*base*_ and *x*_*conflict*_. As such, the data in this figure are by definition noisier, and the analysis has lower statistical power. This leads to smaller effects in the open-loop results compared to main text Figure 3.

Closed-loop stimulation of the right dorsal internal capsule (our most effective open-loop intervention) was more effective than its open loop counterpart at reducing raw RT (the counterpart of *x*_*base*_). Consistent with the specificity illustrated in main text Figure 4C, there was no advantage for closed-loop stimulation on the conflict-specific RT (the counterpart of *x*_*conflict*_). All formatting follows the conventions of main text Figure 4.

**Figure S11:**
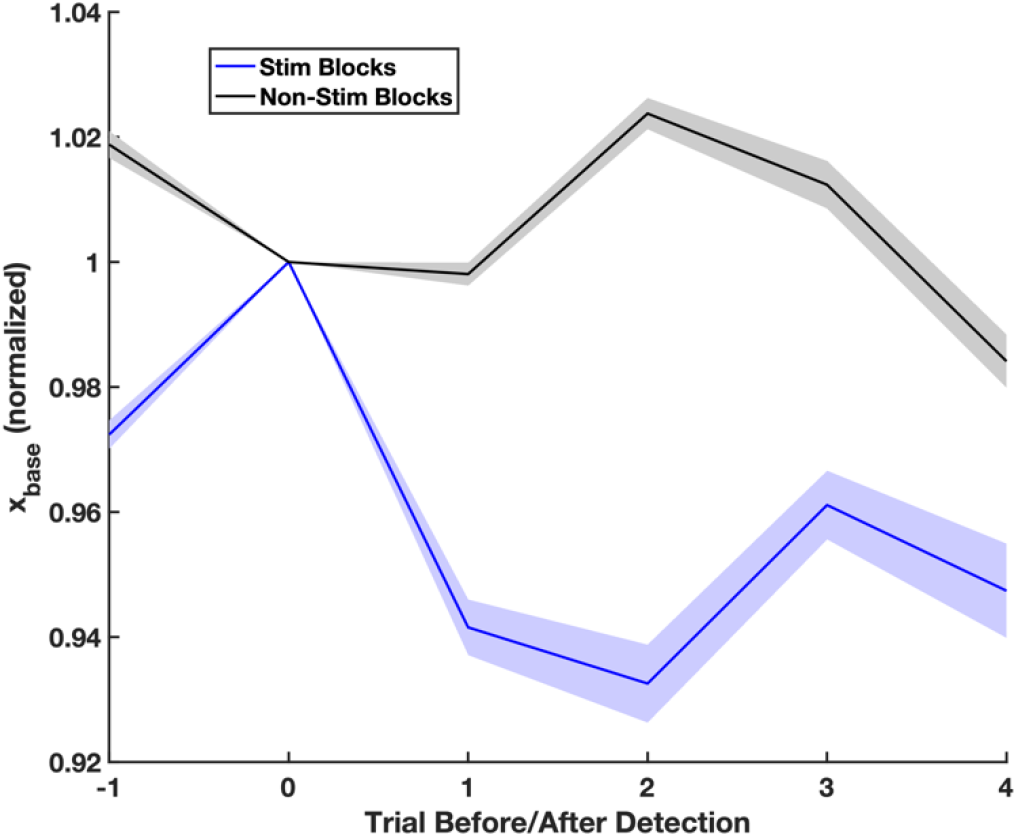
Closed loop stimulation effects cannot be explained by regression to the mean. For the 3 participants included in closed loop experiments, we examined the non-stimulated blocks they performed prior to closed-loop work. (These were the same blocks of behavior data used to train the closed loop model.) We applied the threshold that was used for actual closed loop work in that participant, then detected points in the non-stimulated blocks where our closed loop detector would have triggered (i.e., where *x*_*base*_ was above threshold). We compared these trials to the trials during closed loop stimulation, where the detector actually did trigger and stimulation was delivered. For these two sets of trials, we plot *x*_*base*_ before and after the detection event. The curves show this “stimulation triggered average”, for blocks where stimulation was delivered (blue) or was not (black). The line represents the grand mean over all such events and patients, while shading represents standard error of the mean. To enable averaging over trials and patients (and to highlight changes), we normalized all data such that the detection trial always had a state value of 1.

During active stimulation, *x*_*base*_ decreases (RT improves) on the very next trial, and remains low. In the absence of stimulation, it remains effectively flat, perhaps oscillating around a baseline. This clear difference between trajectories demonstrates that, in the absence of stimulation, our observed performance improvements are highly unlikely.

**Figure S12:**
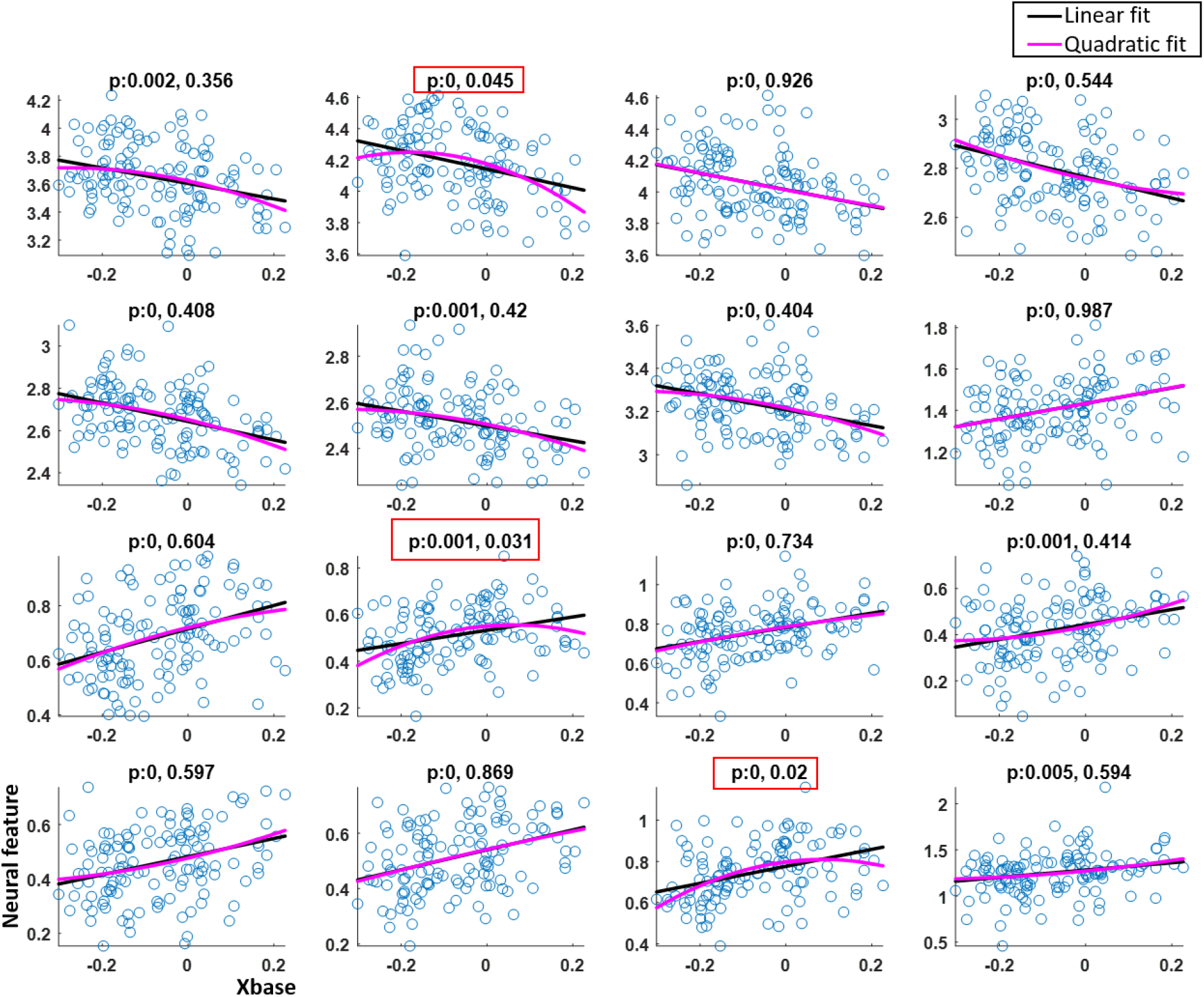
Scatter plot of *x*_*base*_ and neural features in an example dataset. The p-values corresponding to each subplot correspond to the linear and quadratic coefficients in the model Y~1+X and Y~1+X+X^2^ respectively. Most features show a linear relationship (significant linear coefficient only). The features that show both significant linear and quadratic coefficients are boxed in red. In each of those cases, the linear model is still a better fit to the data.

**Figure S13:**
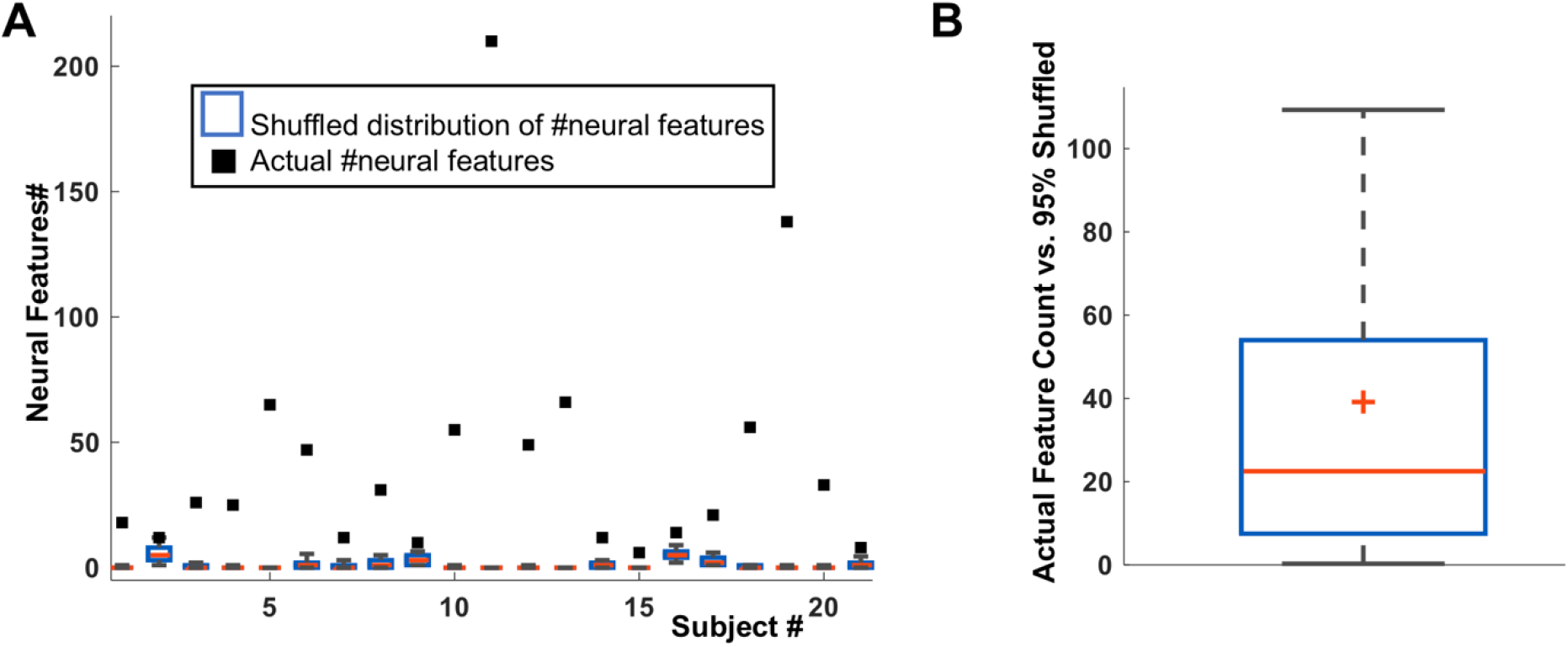
Comparison of neural decoding vs. chance performance. Our neural decoding and variable selection algorithm automatically prunes the feature set to only those LFP power variables that are strongly correlated with the behavioral outcomes (here, *x*_*base*_ and *x*_*conflict*_). As in ref. 70, we tested whether these correlations could occur by chance. We randomly shuffled the order of the behavior trials, then re-ran the decoder variable selection. We repeated this process 100 times for each participant, and on each step, counted the number of features (power bands in specific channels) that were selected. This provides a distribution of the number of significant encoding features expected in purely random data. A), individual-level data. Each black square shows the actual number of features selected by the decoder construction algorithm. The box plot (blue/red; in many cases small due to compression near 0) shows the numbers of features selected by the same algorithm on randomly shuffled data. When the actual number of features is well outside this permutation null distribution, it indicates that the decoder is leveraging actual information present in the dataset. This is visibly true, with a very large separation, for almost all participants in the dataset. B), aggregate data. This boxplot shows the difference between the actual number of features selected (black squares in panel A) and the 95^th^ quantile of the null distribution (top of the blue box in panel A), aggregated across participants. There is a median gap of 20 features, with a confidence interval excluding 0, again demonstrating that the decoding models are capturing true structure in the data.

**Figure S14:**
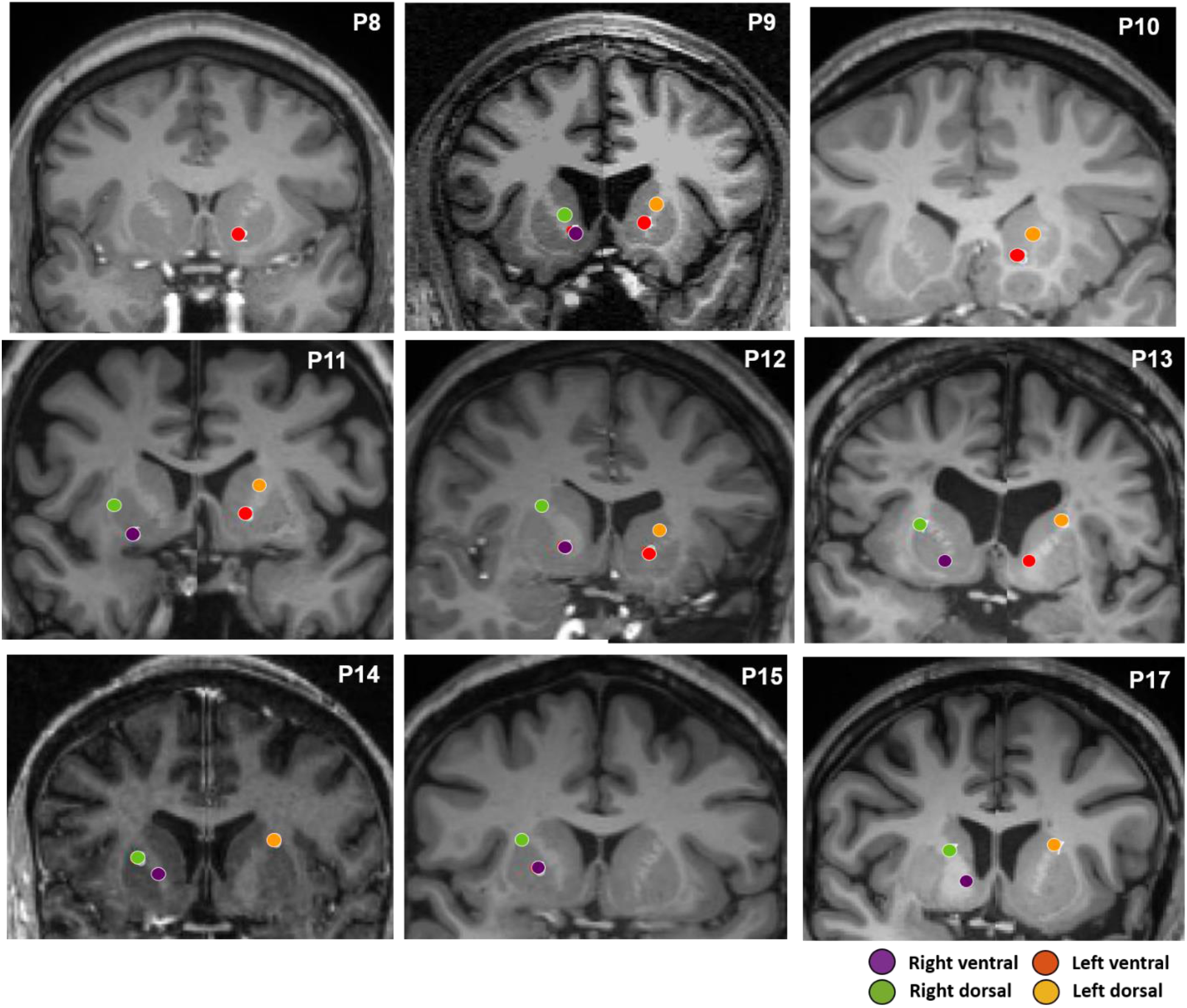
Capsular electrode placements in 9 participants, derived from post-operative CT registered to pre-operative MRI. The last row corresponds to participants who performed the closed loop MSIT experiment, and the first two rows to participants in the open loop experiment.

**Table S1:** Clinical characteristics of participants. Age is rounded to nearest decade to mask participant identity. Seizure focus represents the area ultimately targeted for resection or responsive neurostimulator implant based on clinical consensus after acute monitoring. Neuropsychiatric diagnoses were, when available, taken from neuropsychological testing performed prior to implant. The “Depression Score” is the score on the Beck Depression Inventory (BDI-II). The “Anxiety Score” is the Beck Anxiety Inventory (BAI). In cases where the testing neuropsychologist chose to use a different scale, the name of that scale is given in parentheses. The listed anti-seizure medications are those that were being given at the time of stimulation experiments. Clonazepam is listed either as an anti-seizure or “other” medication depending on whether it was being used to reduce seizure frequency or manage anxiety. ADHD: Attention Deficit Hyperactivity Disorder GAD: Generalized Anxiety Disorder scale PHQ: Personal Health Questionnaire PMDD: Pre-Menstrual Dysphoric Disorder PSWQ: Penn State Worry Questionnaire PTSD: Post-Traumatic Stress Disorder

| P# | Age | Seizure Focus | Prior Neuro-Psychiatric Diagnoses | Depression Score | Anxiety Score | Anti-Seizure Medications | Other CNS Medications |
| --- | --- | --- | --- | --- | --- | --- | --- |
| 1 | 40 | Right mesial temporal | None | 14 | 38 (PSWQ) | Lamotrigine, zonisamide | Clonazepam |
| 2 | 40 | Left inferior temporal/superior temporal | None | 36 | 14 | Oxcarbazepine | Mirtazapine |
| 3 | 30 | Left temporal | None | N/A | N/A | Ezogabine, Levetiracetam | Hydromorphone |
| 4 | 60 | Left insula | Depression, suicidality, visual hallucinations | N/A | N/A | Carbamazepine, levetiracetam, zonisamide | N/A |
| 5 | 20 | Right frontal region and left superior parietal lobule | None | 1 | 1 | Clobazam, cannabidiol, lacosamide, | N/A |
| 6 | 50 | Bitemporal | None | Score not recorded (normal) | Score not recorded (normal) | Levetiracetam | Oxycodone |
| 7 | 20 | Left temporo-occipital | ADHD | N/A | N/A | Lamotrigine, oxcarbazepine, zonisamide | N/A |
| 8 | 20 | Right frontotemporal | "at risk" for attentional disorder | 7 | 4 | Lacosamide, topiramate | N/A |
| 9 | 40 | Multifocal (mainly right temporal) | None | 21 | 15 | Levetiracetam, phenytoin | Clonazepam |
| 10 | 20 | Right mesial temporal | Depression | 8 | 3 | Lacosamide, lamotrigine | Clonazepam |
| 11 | 60 | Left anterior mesial temporal lobe | Depression, insomnia | 25 | 20 | Clobazam, eslicarbazepine, lamotrigine | Hydromorphone, oxycodone |
| 12 | 20 | Left anterior cingulate | Depression | 0 | 3 | Gabapentin, lacosamide, lamotrigine | N/A |
| 13 | 50 | Bitemporal (left more than right) | None | 8 | 2 | Gabapentin, levetiracetam, zonisamide, | Oxycodone |
| 14 | 50 | Bilateral hippocampal | PMDD | 11 | 5 | Clobazam, levetiracetam | N/A |
| 15 | 40 | Left posterior mesial temporal | None | 12 | 12 | Lamotrigine | N/A |
| 16 | 30 | Left temporal | None | 0 | 2 | Lamotrigine, zonisamide | N/A |
| 17 | 40 | Bitemporal | ADHD, depression, anxiety, PTSD, substance abuse | 33 | 32 | Clonazepam, gabapentin, lamotrigine | Acetaminophen-butalbital-caffeine, oxycodone |
| 18 | 40 | Multifocal: left orbitofrontal region, right mesial frontal, right lateral temporal, right middle temporal | None | N/A | N/A | Lamotrigine, valproate | N/A |
| 19 | 30 | Right frontotemporal parietal | Bulimia nervosa, depression | 2 | 4 | Clobazam, topiramate | N/A |
| 20 | 30 | Multifocal: primarily right posterior | Depression, suicidality | 27 (PHQ-9) | 20 (GAD-7) | Lamotrigine, levetiracetam | N/A |
| 21 | 40 | Bilateral mesial temporal | None | 5 | 1 | Lamotrigine, zonisamide | N/A |

**Table S2:** Order of stimulation blocks in participants who received open-loop and closed loop (CL) capsular stimulation. (DC: Dorsal Capsule, VC: Ventral Capsule, L:Left, R:Right)

| P# | 8 | 9 | 10 | 11 | 12 | 13 | 14 (CL) | 15 (CL) | 17 (CL) |
| --- | --- | --- | --- | --- | --- | --- | --- | --- | --- |
| Blocks | L VC | R VC<br>L DC<br>R DC<br>L VC | L DC<br>L VC<br>L DC | R VC<br>R DC<br>L VC<br>L DC | R VC<br>R DC<br>L VC<br>L DC<br>R DC | R VC<br>R DC<br>L VC<br>L DC | R DC<br>L DC<br>L DC<br>R VC<br>R DC | R DC<br>R DC<br>R VC | R DC<br>R DC<br>L DC<br>R VC<br>R DC<br>R DC<br>R VC |

**Table S3a:**
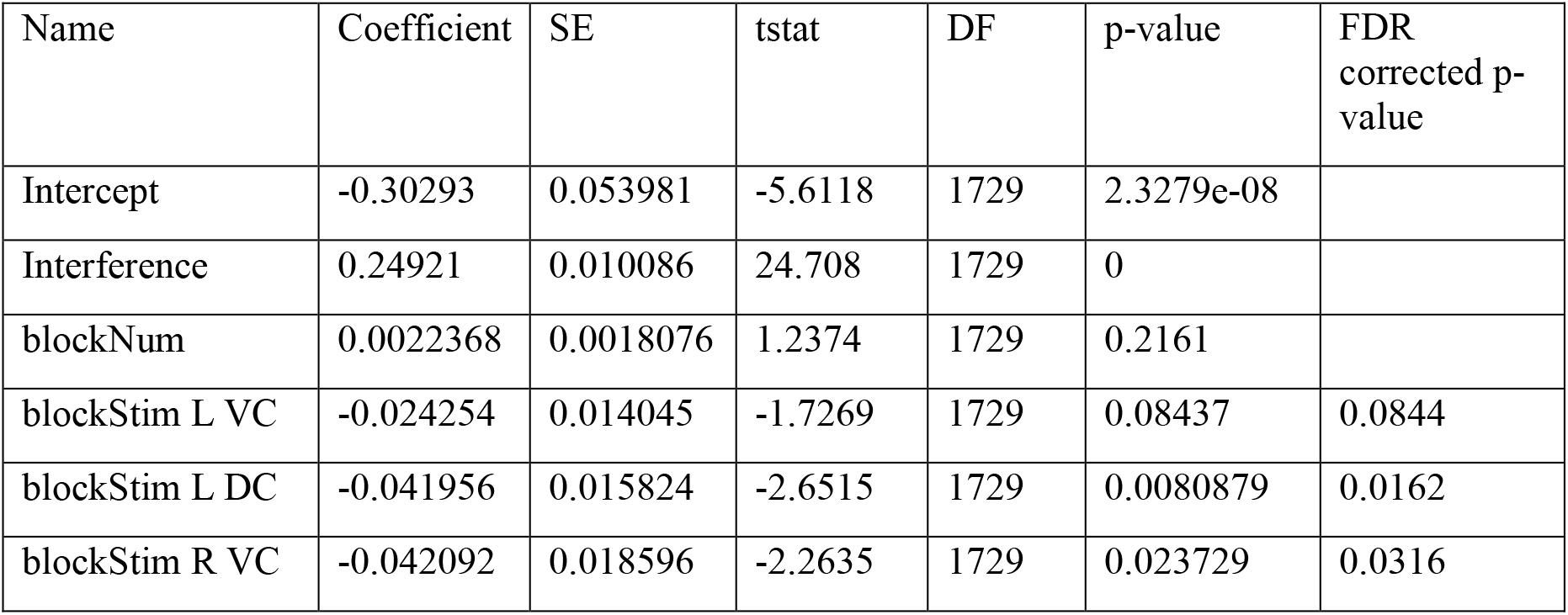
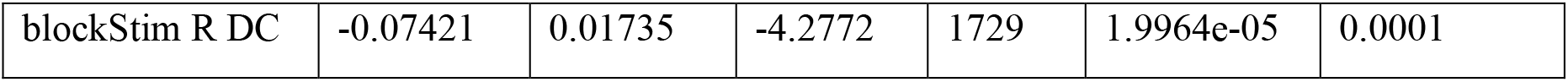
Regression coefficients for reaction time (RT) in the open-loop stimulation experiments, from a generalized linear mixed-effects model, with a log-normal distribution and identity link function. Formula: RT ~ Conflict + blockNum + blockStim + (1 | Participant) Fixed effects coefficients (95% CIs):

**Table S3b:**
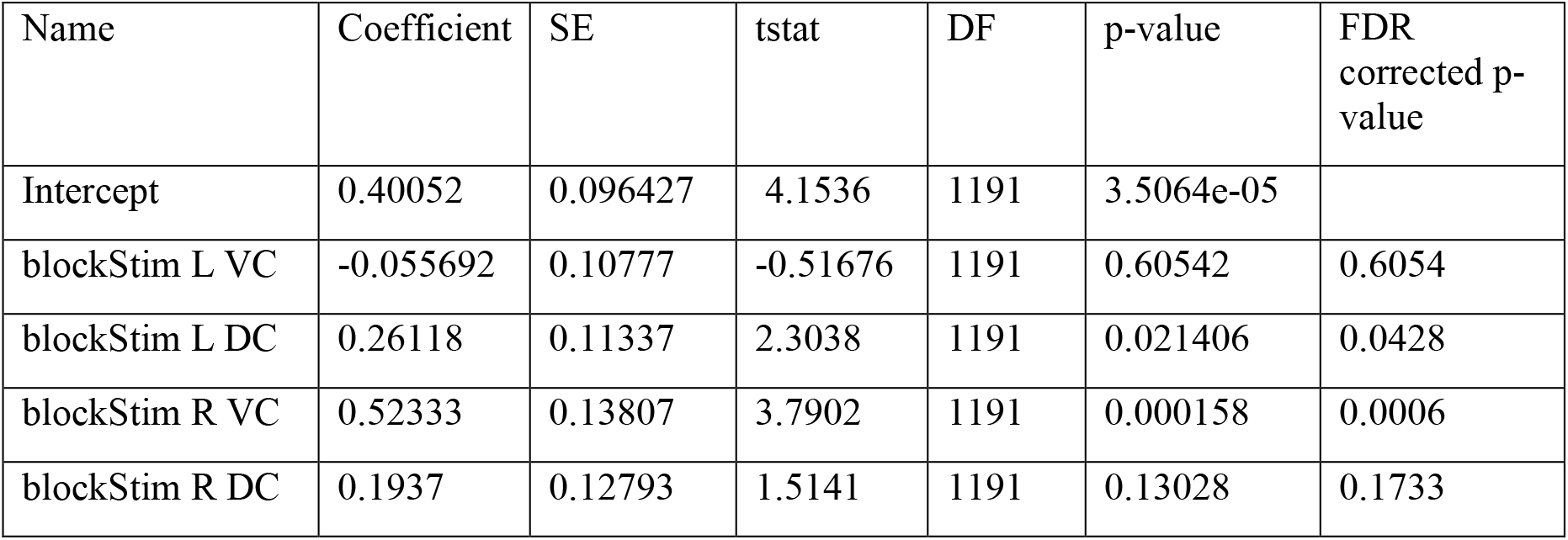
Regression coefficients for theta power in the open-loop stimulation experiments, from a generalized linear mixed-effects model, with a log-normal distribution and identity link function. Formula: Theta ~ blockStim + (1 | Participant) Fixed effects coefficients (95% CIs):

| Name | Coefficient | SE | tstat | DF | p-value | FDR corrected p-value |
| --- | --- | --- | --- | --- | --- | --- |
| Intercept | 0.40052 | 0.096427 | 4.1536 | 1191 | 3.5064e-05 |  |
| blockStim L VC | -0.055692 | 0.10777 | -0.51676 | 1191 | 0.60542 | 0.6054 |
| blockStim L DC | 0.26118 | 0.11337 | 2.3038 | 1191 | 0.021406 | 0.0428 |
| blockStim R VC | 0.52333 | 0.13807 | 3.7902 | 1191 | 0.000158 | 0.0006 |
| blockStim R DC | 0.1937 | 0.12793 | 1.5141 | 1191 | 0.13028 | 0.1733 |

**Table S4:**
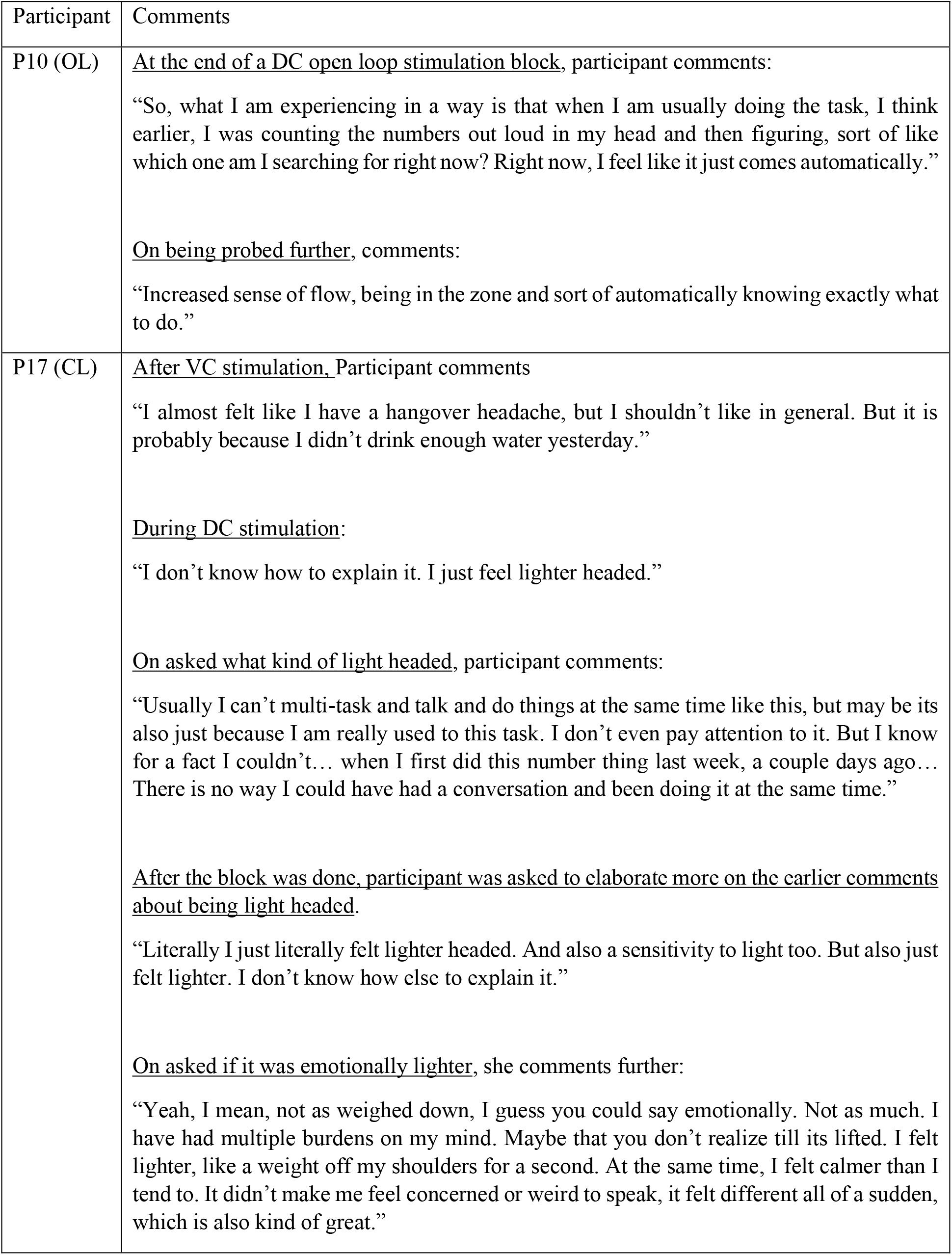
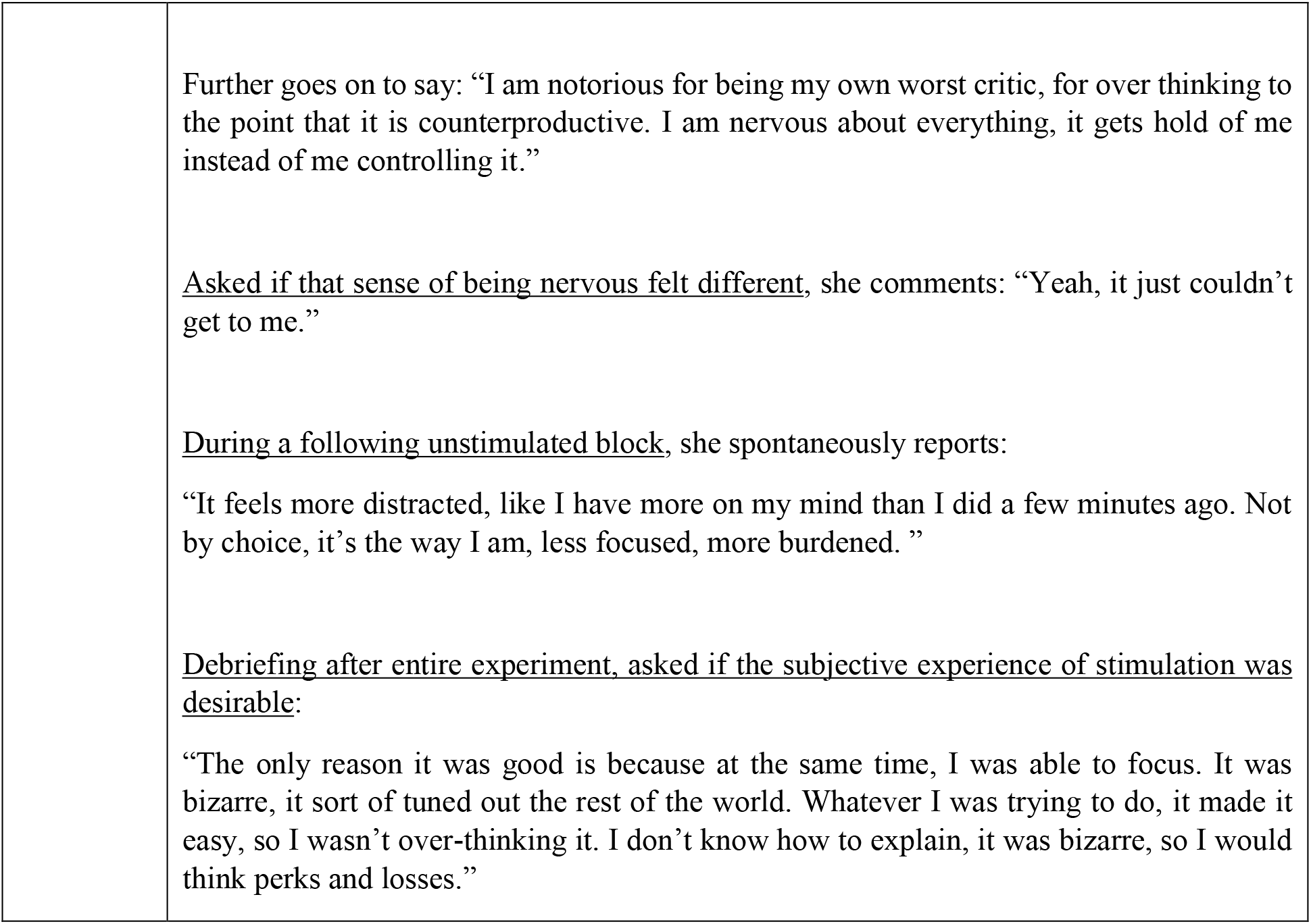
Qualitative comments made by 2 participants during and after stimulated MSIT sessions, demonstrating relief of anxiety and greater ability to redirect attention.

**Table S5:**
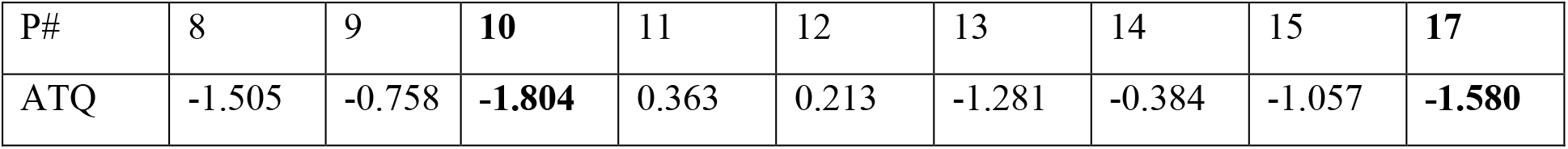
Self-report of positive emotional effects from capsular neurostimulation compared with prior self-report of self-control. As part of a broader study in which these experiments were embedded (Widge et al., 2017), participants completed a variety of self-report measures. These were compared against a companion dataset of 36 healthy controls, and are expressed as Z-scores for each participant relative to the healthy population. The Effortful Control subscale of the Adult Temperament Questionnaire (ATQ, 77-item short form) reports a respondent’s capability to focus attention and shift behavior when appropriate. The two participants in Table S3 who reported subjective positive effects of stimulation are highlighted in bold. They also had the most negative (impaired) scores on ATQ Effortful Control. We did screen other questionnaires and did not observe this pattern; this was not a pre-planned analysis.

| P# | 8 | 9 | **10** | 11 | 12 | 13 | 14 | 15 | **17** |
| --- | --- | --- | --- | --- | --- | --- | --- | --- | --- |
| ATQ | -1.505 | -0.758 | **-1.804** | 0.363 | 0.213 | -1.281 | -0.384 | -1.057 | **-1.580** |

**Table S6:** Number of trials used for training, validation and testing of the encoder-decoder model in each participant. Note that the Train/Validate/Test columns will add to more than the overall number of trials, because there was overlap between Train and Validate sets. The Test set was fully disjoint from both Train and Validate. For P3, an encoder model could not be estimated for *x*_*conflict*_ even when using 70% of the available trials.

| P# | Total Trials | Baseline |  |  | Conflict |  |  |
| --- | --- | --- | --- | --- | --- | --- | --- |
|  |  | Train | Validate | Test | Train | Validate | Test |
| 1 | 192 | 128 | 80 | 48 | 144 | 83 | 35 |
| 2 | 383 | 153 | 114 | 114 | 194 | 137 | 160 |
| 3 | 304 | 153 | 126 | 101 | NA | NA | NA |
| 4 | 304 | 152 | 114 | 114 | 205 | 128 | 70 |
| 5 | 250 | 166 | 111 | 63 | 166 | 111 | 63 |
| 6 | 304 | 204 | 135 | 75 | 204 | 135 | 75 |
| 7 | 304 | 204 | 135 | 75 | 204 | 135 | 75 |
| 8 | 312 | 159 | 111 | 94 | 132 | 111 | 135 |
| 9 | 305 | 169 | 118 | 102 | 169 | 118 | 102 |
| 10 | 250 | 117 | 69 | 66 | 175 | 54 | 64 |
| 11 | 223 | 141 | 80 | 72 | 141 | 80 | 72 |
| 12 | 310 | 144 | 95 | 143 | 144 | 95 | 143 |
| 13 | 337 | 213 | 130 | 100 | 165 | 106 | 135 |
| 14 | 287 | 140 | 114 | 103 | 140 | 114 | 103 |
| 15 | 241 | 105 | 97 | 91 | 105 | 97 | 91 |
| 16 | 394 | 178 | 197 | 108 | 178 | 197 | 108 |
| 17 | 296 | 110 | 144 | 86 | 110 | 144 | 86 |
| 18 | 270 | 170 | 118 | 75 | 180 | 120 | 67 |
| 19 | 258 | 172 | 114 | 64 | 170 | 113 | 63 |
| 20 | 444 | 256 | 158 | 94 | 128 | 96 | 96 |
| 21 | 372 | 184 | 165 | 115 | 200 | 284 | 284 |

